# Transferable deep generative modeling of intrinsically disordered protein conformations

**DOI:** 10.1101/2024.02.08.579522

**Authors:** Giacomo Janson, Michael Feig

**Affiliations:** Department of Biochemistry and Molecular Biology, Michigan State University, East Lansing, Michigan, USA

## Abstract

Intrinsically disordered proteins have dynamic structures through which they play key biological roles. The elucidation of their conformational ensembles is a challenging problem requiring an integrated use of computational and experimental methods. Molecular simulations are a valuable computational strategy for constructing structural ensembles of disordered proteins but are highly resource-intensive. Recently, machine learning approaches based on deep generative models that learn from simulation data have emerged as an efficient alternative for generating structural ensembles. However, such methods currently suffer from limited transferability when modeling sequences and conformations absent in the training data. Here, we develop a novel generative model that achieves high levels of transferability for intrinsically disordered protein ensembles. The approach, named idpSAM, is a latent diffusion model based on transformer neural networks. It combines an autoencoder to learn a representation of protein geometry and a diffusion model to sample novel conformations in the encoded space. IdpSAM was trained on a large dataset of simulations of disordered protein regions performed with the ABSINTH implicit solvent model. Thanks to the expressiveness of its neural networks and its training stability, idpSAM faithfully captures 3D structural ensembles of test sequences with no similarity in the training set. Our study also demonstrates the potential for generating full conformational ensembles from datasets with limited sampling and underscores the importance of training set size for generalization. We believe that idpSAM represents a significant progress in transferable protein ensemble modeling through machine learning.

**AUTHOR SUMMARY:** Proteins are essential molecules in living organisms and some of them have highly dynamical structures, which makes understanding their biological roles challenging. Disordered proteins can be studied through a combination of computer simulations and experiments. Computer simulations are often resource-intensive. Recently, machine learning has been used to make this process more efficient. The strategy is to learn from previous simulations to model the heterogenous conformations of proteins. However, such methods still suffer from poor transferability, meaning that they tend to make incorrect predictions on proteins not seen in training data. In this study, we present idpSAM, a method based on generative artificial intelligence for modeling the structures of disordered proteins. The model was trained using a vast dataset and, thanks to its architecture and training procedure, it performs well on not just proteins in the training set but achieves high levels transferability to proteins unseen in training. This advancement is a step forward in modeling biologically relevant disordered proteins. It shows how the combination of generative modeling and large training sets and can aid us understand how dynamical proteins behave.

## INTRODUCTION

A major paradigm of structural biology is that the function of a protein is determined by the properties of its structural ensemble. The level of conformational variability of different proteins can range from relatively rigid molecules to fully unstructured ones[1]. For more rigid proteins, a single three-dimensional (3D) structure, obtainable by experimental[2] or computational prediction methods[3, 4], often provides key insights into its function. On the other hand, for highly dynamic proteins, function may only be fully understood by knowing the statistical properties of their structural ensembles[5, 6]. Intrinsically disordered proteins (IDPs) or intrinsically disordered regions (IDRs) of otherwise more rigid proteins exhibit very high levels of conformational flexibility[7]. For simplicity, we will use the term IDR to refer to both in the remainder of the text. IDRs have many important biological roles and perfectly exemplify the difficulty of investigating structure-to-function relationships for dynamic proteins[8]. Such studies require the integration of experimental data and computational methods for generating pools of 3D structures compatible with that data[9]. Therefore, algorithms for modeling conformational ensembles are central for our understanding of a large and biomedically relevant fraction of the protein space.

A powerful method to generate structural ensembles consists of physics-inspired sampling strategies, such as molecular dynamics[10] (MD) or Markov Chain Monte Carlo[11] (MCMC) simulations. While there has been progress in improving the force fields to model protein dynamics in atomistic MD and especially for IDRs[12], the high computational cost of running such simulations still limits their usage even on the most advanced current computing hardware. IDRs are particularly challenging because the more extended and dynamic conformations require both larger simulation systems and longer simulation times to generate representative conformational ensembles. Coarse-grained[13] or implicit solvent[14] simulations can reduce the cost but at the expense of introducing modeling approximations.

Recently, an alternative approach has emerged in the form of machine learning algorithms[15-17]. The general idea is to train a machine learning model on molecular simulation data to learn to sample new conformations in a computationally efficient way[18]. Deep generative models are a promising strategy for achieving this goal thanks to their capability of modeling complex probability distributions and fast sampling with one or few steps[19-21]. They have been applied to datasets of both folded[22] and intrinsically disordered proteins or peptides[23]. Most applications focus on replicating ensembles from simulations, but recently the DynamICE model[24] was used to integrate experimental data in the ensemble generation process of IDRs, highlighting the potential of generative models in integrative structural biology[9]. In closely related work, generative models such as RFdiffusion[25] and FrameDiff[26] have also been applied successfully in protein design studies[27], where they are trained on experimentally-determined folded proteins instead of molecular simulation datasets.

Despite these advancements, the problem of rapidly generating a useful conformational ensemble for an arbitrary dynamic protein remains essentially an unsolved problem. While nowadays it is routinely possible to train system-specific models that may recapitulate or even extrapolate sampling seen in training data, the critical challenge is to obtain transferable models that can be applied to new systems with satisfactory performance[28], especially for complex systems such as IDRs[29]. The fundamental issue is that deep learning methods are highly data-driven, whereas transferability requires generalization beyond given data in order to model previously unseen systems. General principles could be introduced via physics-based terms, following the typical approach in traditional simulations, but additional sampling requirements may negate the advantages of direct ensemble generation via deep generative models. Alternatively, a very large, diverse training set may provide enough information to learn general principles with a sufficiently deep network to reach transferability.

Earlier efforts by us towards generating ensembles for intrinsically disordered peptides in a transferable manner have resulted in idpGAN[29], based on a generative adversarial network[20] (GAN). Two versions of idpGAN were trained, one based on the residue-based coarse-grained model COCOMO[13], another one on sampling with the ABSINTH implicit solvent potential[14, 30]. For COCOMO-based sampling we achieved good transferability for most test proteins, but since COCOMO focuses on polymer-like properties and is limited in capturing complex conformational distributions, the generated ensembles by idpGAN are equally limited in capturing detailed aspects of IDRs. The idpGAN version trained on ABSINTH simulations had the potential to generate more realistic conformational distributions for IDRs, but we found that transferability was only achieved for some test proteins, therefore limiting its usefulness in broader applications.

Here we report idpSAM (Structural Autoencoder generative Model) that significantly improves over our previous efforts. IdpSAM achieves fully transferable ensemble generation for IDRs based on the ABSINTH model. The key factors are a new type of generative model and significantly expanded training data.

We switched from the GAN model used earlier to a denoising diffusion probabilistic model[19] (DDPM) motivated by the recent success of such models in other recently proposed generative frameworks. Diffusion models can be directly applied to 3D atomic coordinates. However, training such models on protein atomic coordinates in a transferable way has been shown to be challenging, for example requiring large pre-trained networks[25] and bespoke geometrical-aware architectures. For this reason, we explore here an alternative approach and make use of latent DDPMs, which operate on a learned “latent” representation of the original data. Our approach has been inspired by latent DDPMs employed for image generation[31-33]. Recently, GEOLDM, a model of this kind, has been applied to generate drug-like molecules obtaining state-of-the-art performance[34]. A similar approach was also independently adopted in protein design studies[35]. Our method differs from these models as we introduce the use of Euclidean-invariant encodings. This choice enables the incorporation of any expressive neural network into the DDPM, such as the commonly used transformers[36], instead of being limited to only E(3)-equivariant networks. To our knowledge, this is the first latent DDPM to have been applied to datasets consisting of protein simulations. The method is also different from a previously-reported work for sampling IDR conformations via the latent space of an autoencoder[37], since that approach was non-transferable and did not involve a generative model component to learn the structure of the latent space.

Along with the improved generative model, we also significantly expanded the underlying training set. We decided to continue to work with ABSINTH simulations because, despite their approximations in the treatment of solvent, proteins are represented in atomistic details and, more importantly, because ABSINTH captures non-trivial sequence-specific interaction patterns that result in complex distributions of conformations involving the formation of realistic transient secondary structure elements[38, 39]. Yet, ABSINTH is still computationally efficient enough to generate very large amounts of training data. It would otherwise be very difficult to achieve such data via fully explicit solvent atomistic simulations, especially for IDRs. The increased amount of training data also allowed us to address questions of data sufficiency for training transferable ensemble generating models that have been largely unexplored to date.

As in our previous work, we train a system to model only the Cα traces of IDRs. This made the machine learning effort manageable with available hardware. However, we can recover full atomistic detail rapidly and accurately with the cg2all method developed recently[40]. This allows us to introduce a new platform for rapidly generating IDR ensembles at different resolutions via a deep learning framework that can potentially replace traditional simulation approaches.

## RESULTS

### Structural Autoencoder Model

In order to generate conformations of IDRs, a structural autoencoder model (SAM) was trained with ABSINTH simulation data. An overview of the training and sampling processes of idpSAM are shown in **Fig 1**. SAM consists of two main components: first, an autoencoder (AE) is trained to generate SE(3)-invariant encodings of Cα coordinates (**Fig 1A**). Second, a denoising diffusion probabilistic model (DDPM) is employed to learn the probability distribution of encodings from the encoder network with amino acid sequences as conditional input (**Fig 1B**). Once both systems have been trained, sampling of conformations for a given peptide sequence is accomplished by generating encodings via the diffusion model and converting them to 3D structures with the decoder network of the AE (**Fig 1C**).

**Fig 1.**
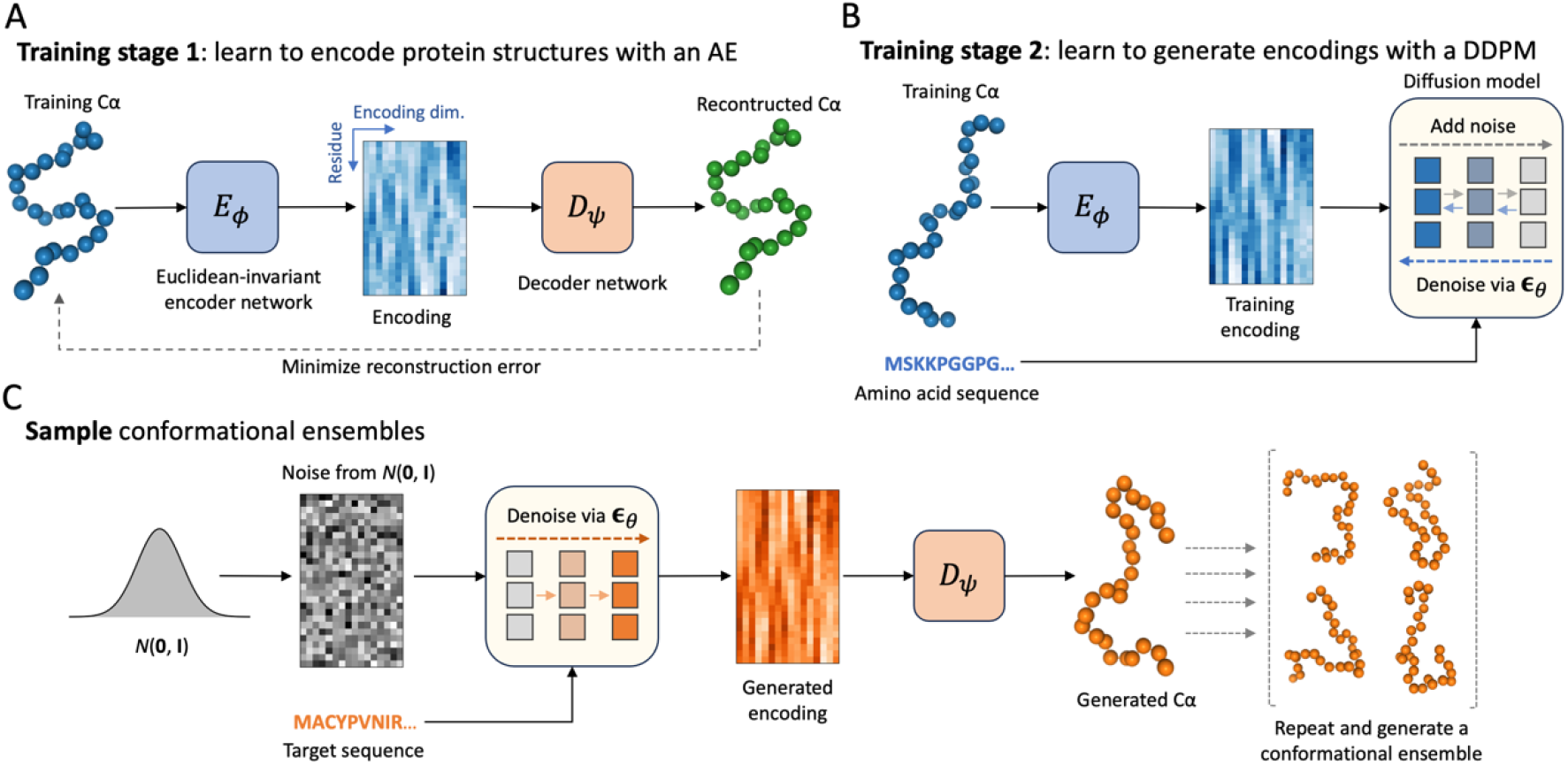
Architecture of the idpSAM generative model. (A) and (B) Two-staged training process of the structural autoencoder model (SAM). In the first stage, a Euclidean-invariant AE is trained to encode Cα protein conformations. An example Cα structure from the training set is shown along with a 2D image representation of its encoding. In the image, the horizontal axis represents the encoding dimension (encodings have *c* = 16 channels) and the vertical axis represents the residue index. In the second stage, a DDPM is trained to learn the distribution of the encodings in the training set. (C) Generating samples via SAM. Once the DDPM is trained, it can be used to generate encodings for new peptide sequences via reverse diffusion starting from Gaussian random noise. The decoder then maps the encodings to 3D structures. The process can be repeated to generate full conformational ensembles.

Since the DDPM of SAM operates over encoded representations of 3D conformations, the performance of the decoder is a crucial component for generating 3D structures. To assess the reconstruction accuracy of the decoder, we evaluated it on MCMC snapshots of test set peptides encoded via the encoder network. In **S1 Fig** we show examples of how the decoder can correctly recover Cα-Cα distances and α torsion angles of test encoded conformations. Quantitative analyses are reported in **S1 Table**. As a control, we computed the *C*_*dist*_ and *C*_*tors*_ values (see Methods) for test set conformations with perturbed copies of themselves, generated by adding Gaussian noise with *σ* values ranging from 0.01 to 0.1 Å. The average *C*_*dist*_ and *C*_*tors*_ values obtained via reconstruction with the AE are lower than the control performed with noise as little as *σ* = 0.1 Å, indicating that the reconstruction quality is consistently high.

### Transferable modeling of Cα structural ensembles

The goal of idpSAM as a transferable model is to generate 3D structural ensembles for sequences not present in the training set. To evaluate its transferability, we used its DDPM and decoder to generate ensembles for test set peptides. Properties of the generated ensembles for 5 selected peptides are reported in **Fig 2**. The remaining peptides are shown in **S2 Fig** and **S3 Fig**.

**Fig 2.**
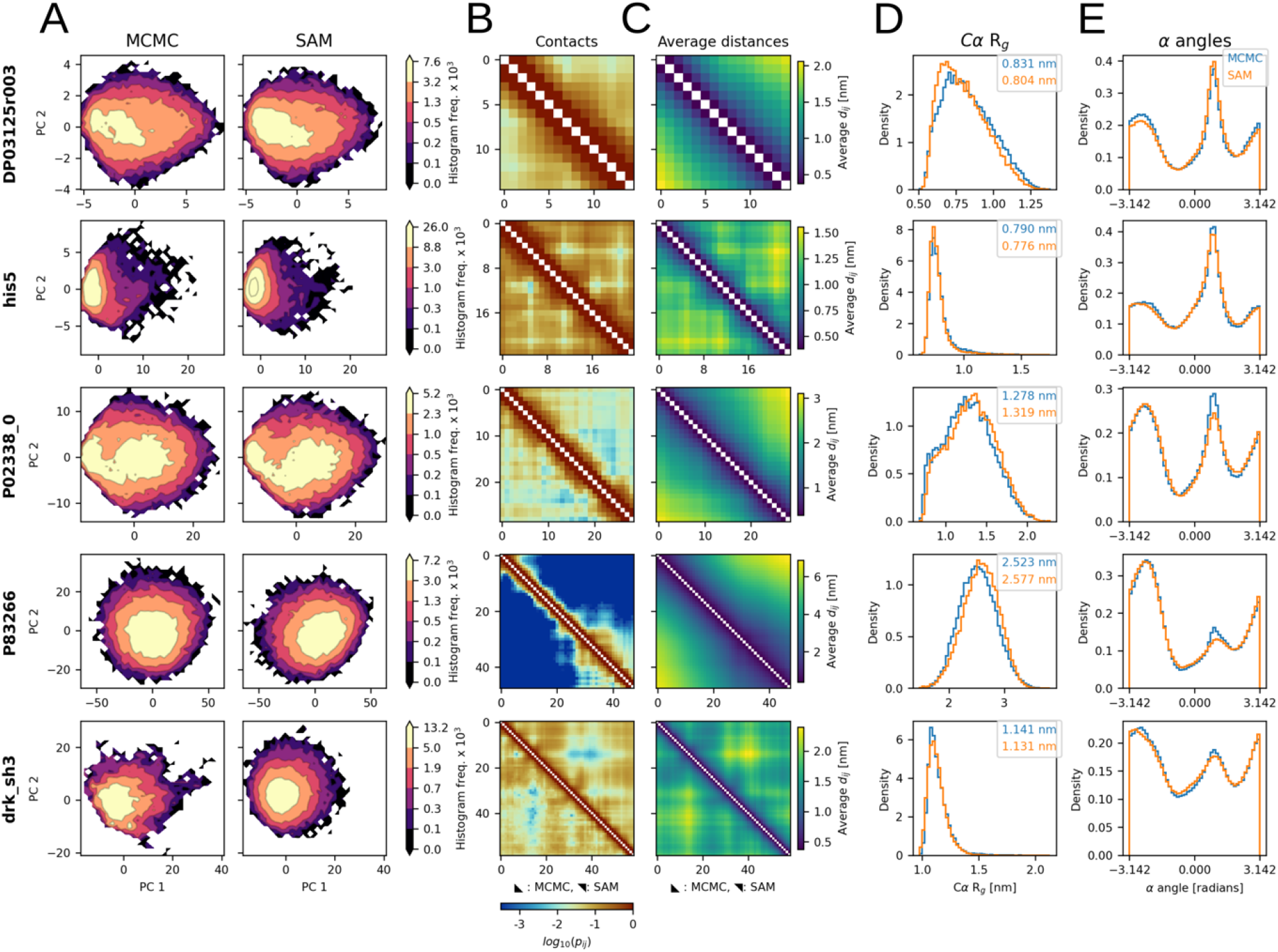
Structural ensembles from idpSAM. Each row shows the MCMC and SAM ensembles of a peptide from the test set: DP03125r003 (*L* = 15), his5 (*L* = 24), P02338_0 (*L* = 29), P83266 (*L* = 48) and drk_sh3 (*L* = 59). (A) Histograms of MCMC (left) and SAM (right) conformations projected along PCA coordinates. Frequency values in colorbars are multiplied by 1 × 10^3^. (B) Cα-Cα contact maps for the MCMC and SAM ensembles. Each cell represents a residue pair and is colored according to its log10(*pij*) value, where *pij* is the probability of observing a contact between residues *i* and *j* in an ensemble. The contact threshold is 8.0 Å. (C) Average Cα-Cα distance maps from MCMC and SAM. (D) Cα radius-of-gyration histograms from MCMC (blue) and SAM (orange). The average values are reported in the legend. (E) Histograms of all α torsion angle values for four subsequent Cα atoms in a peptide backbone from MCMC and SAM. For (B) and (C) the MCMC and SAM ensembles are shown in the lower and upper triangles respectively.

Different amino acid sequences result in reference MCMC ensembles with sequence-specific patterns. IdpSAM closely reproduces various features for almost all peptides. Close agreement is found for the probability distributions projected onto PCA space (**Fig 2A**), sequence-specific patterns of residue-residue interactions, such as contact maps and average distances of Cα atoms (**Fig 2B** and **C**), chain compactness, as shown by the histograms of radius-of-gyration of Cα atoms (**Fig 2D**), and the preferences in local backbone geometry, based on α torsion angles formed by four consecutive Cα atoms along the peptide chain (**Fig 2E**). Significant deviations are found only in a few cases, and in those, the sampling with idpSAM is qualitatively still similar to the reference ensembles. Since none of the test set peptides, or peptides with similar sequences, were present in the training set (see Methods) the ability to predict their conformational ensembles in close agreement with the MCMC results demonstrates transferability.

### Comparison with idpGAN

In a previous study, we trained the idpGAN model[29] on data for 1089 peptides contained in the full training set used here. To determine whether the latent DDPM of SAM has an algorithmic advance over the GAN-based model, we compared their performances. Since idpGAN has been trained on fewer IDRs, we also re-trained idpSAM (both its AE and DDPM) only on those same IDRs. Average evaluation scores (see Methods) are shown in **Table 1**. Pairwise comparisons for all peptides are shown in **S4 Fig A**. It is clear that SAM provides significantly better approximations of the MCMC ensembles than idpGAN, even when SAM is trained with the same smaller training set as idpGAN. The performance increases further when SAM is trained with an expanded training set (**Table 1** and **S4 Fig B**). We note that we were unable to train idpGAN using the expanded dataset, as we could not stabilize the fragile training process of the GAN[41].

**Table 1.** Evaluation scores of ensembles from SAM and other methods.

| Strategy | Training IDRs<sup>b</sup> | MSE_c | MSE_d [nm<sup>2</sup>] | aKLD_d | KLD_r | aKLD_t |
| --- | --- | --- | --- | --- | --- | --- |
| idpGAN | 1,089 | 4.52 ± 1.57 | 0.08 ± 0.02 | 0.29 ± 0.06 | 0.67 ± 0.14 | 0.14 ± 0.05 |
| idpSAM | 1,089 | 3.30 ± 1.18 | 0.04 ± 0.01 | 0.14 ± 0.02 | 0.20 ± 0.03 | 0.09 ± 0.02 |
| idpSAM | 3,259 | 1.42 ± 0.5 | 0.01 ± 0.006 | 0.06 ± 0.02 | 0.05 ± 0.02 | 0.05 ± 0.02 |
| idpSAM-G<sup>b</sup> | 3,259 | 1.48 ± 0.52 | 0.01 ± 0.004 | 0.05 ± 0.01 | 0.05 ± 0.02 | 0.04 ± 0.01 |
| idpSAM-E<sup>c</sup> | 3,259 | 1.52 ± 0.61 | 0.01 ± 0.003 | 0.05 ± 0.02 | 0.05 ± 0.02 | 0.04 ± 0.01 |
| MCMC-5<sup>d</sup> | - | 7.79 ± 2.8 | 0.02 ± 0.008 | 0.14 ± 0.03 | 0.10 ± 0.03 | 0.16 ± 0.04 |

Average scores are reported along with standard errors for the 22 test set peptides.
<sup>a</sup>Number of peptides used in the training set.
<sup>b</sup>Larger SAM model with 20 transformer blocks (instead of 16).
<sup>c</sup>SAM model with FrameDiff edge update operations.
<sup>d</sup>Ensemble generated by randomly extracting 10,000 conformations from only 5 MCMC simulations at 298 K of a test peptide.

In **S5 Fig**, we compare idpSAM and idpGAN ensembles for Q2KXY0, a test peptide for which idpGAN was not performing well in our previous study. We hypothesized that the poor performance of idpGAN for Q2KXY0 was caused primarily by insufficient training data. We find that idpSAM provides a better approximation already with the same training set, but the ensemble is further improved when the training set is expanded. Therefore, we conclude that both, the improved generative model and an increased amount of training data, benefitted the performance.

### Sampling speed

The sampling speed of a molecular conformation generator is the key to its potential usefulness. DDPMs typically have slower sampling than GANs, because GANs need only one forward pass to generate a sample, while DDPMs need multiple passes, in the order of 100-1000. To assess the performance vs. speed tradeoff for idpSAM, we used the SAM version trained on the full training set and sampled with different numbers of diffusion steps[42]. In **Fig 3**, we show the average evaluation scores for test set ensembles with 10,000 conformations as a function of the average wall-clock time needed to generate them. Even with as little as 10 steps, SAM obtains on average better performance than idpGAN. The modeling accuracy of SAM does not seem to substantially change when using 50 or more steps. We found 100 steps to be a good tradeoff between speed and accuracy for Cα ensemble modeling. With 100 steps, the average GPU wall clock time for generating a test set ensemble with 10,000 conformations is about 4 min. For idpGAN it is 0.8 s, but the greater speed results in greatly reduced accuracy. For comparison, generating an ensemble by running 5 MCMC simulations took on average 509 CPU hours on an Intel Xeon Gold 6248R CPU at 3.00GHz (CAMPARI, the software for running training MCMC simulations, is a CPU-only package).

**Fig 3.**
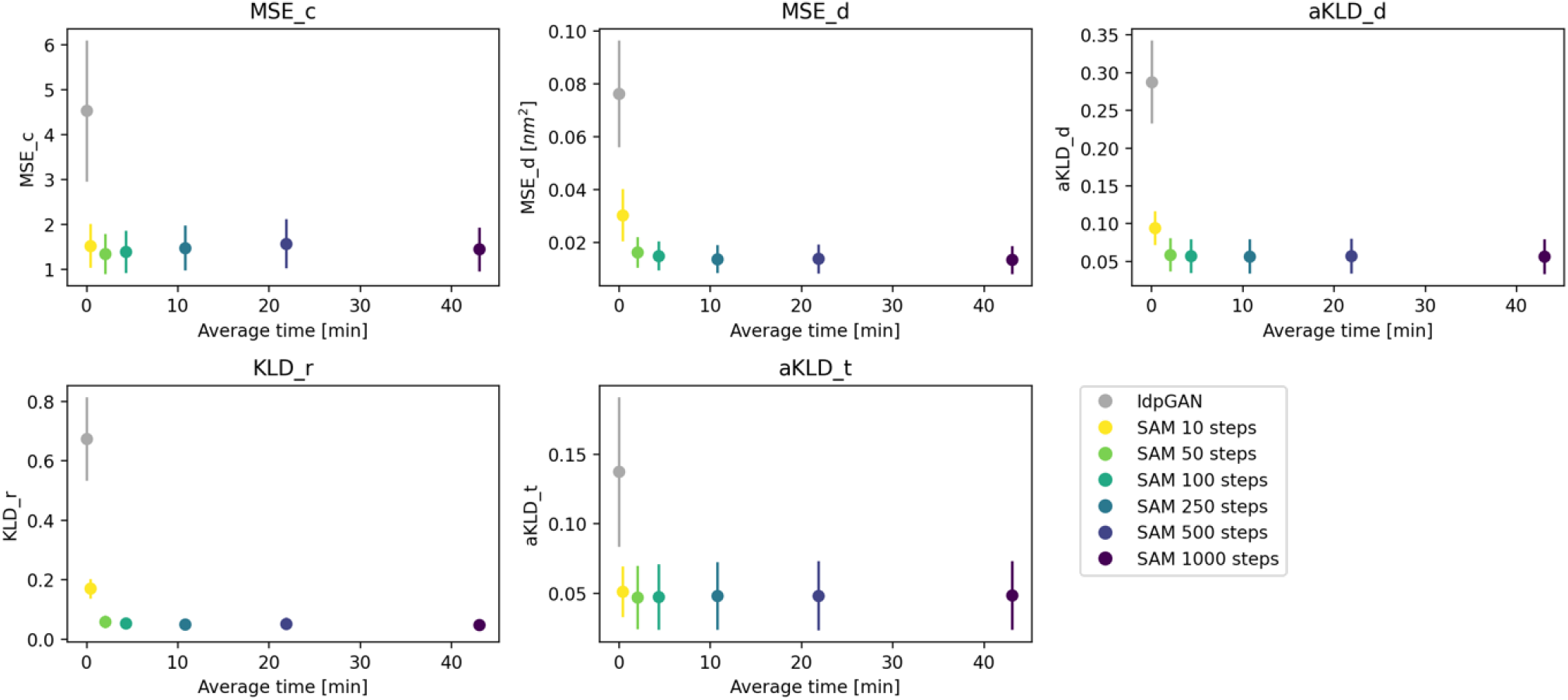
Sampling speed of SAM. Average evaluation scores for the 22 test peptides as a function of the average wall-clock times used to generate their ensembles. Error bars represent the standard error of the mean across peptides. Ensembles of 10,000 structures were generated by running models implemented with PyTorch on a NVIDIA RTX2080Ti GPU. A batch size of 256 was used for all models. Each subplot reports a different score.

### Case studies illustrating sequence and structure transferability

In the test set, we included a pair of similar sequences: the wild-type (wt) sequence and a designed mutant[43] of the calcitonin gene-related peptide[44] (CGRP). The two 37-residue sequences differ by only 4 amino acid substitutions (**S2 Table**). We used them to assess the capability of SAM to capture the effect of sequence variations. The reference MCMC ensembles of the wild-type and mutant peptides share similarities, yet they have some clear distinctions. For example, differences in contact and average Cα-Cα distance maps show that the C-terminal region (residues 25-37) behaves differently in the two peptides. SAM appears to correctly capture such variations (**Fig 4A** and **B**).

**Fig 4.**
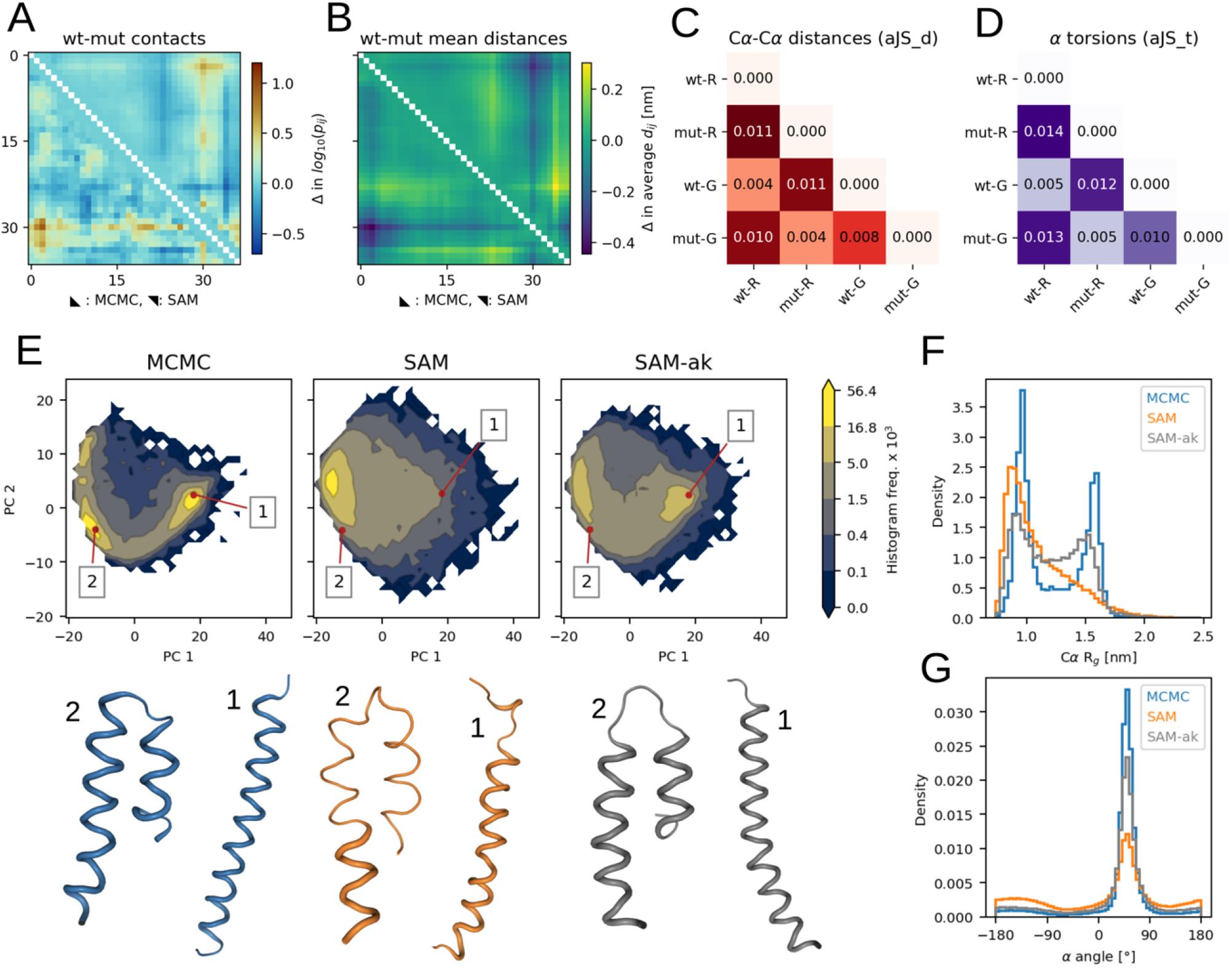
Two case studies from the test set. (A) to (D) data for CGRP peptides. (E) to (G) data for the ak37 peptide. (A) Differences in Cα-Cα contact log10(*pij*) between the wt and mutant CGRP ensembles. Values from reference MCMC ensembles are in the lower triangle, values from SAM are in the upper one. (B) Differences in average Cα-Cα distances between the wt and mutant CGRP ensembles. (C) aJSD_d scores for the following CGRP ensembles: wt and mutant from reference MCMC simulations (wt-R and mut-R, respectively) and wt and mutant ensembles generated by SAM (wt-G and mut-G). (D) aJS_t scores considering α angle distributions. (E) PCA histograms for ak37 ensembles from: MCMC (left), default SAM (center) and a SAM version in which three ak peptides were added for training (SAM-ak, right). Frequency values are multiplied by 1 × 10^3^. For each histogram, two snapshots extracted from the highlighted points are shown below. These snapshots belong to fully-helical (1) or helical hairpin (2) states. (F) Histograms of Cα *R*_g_ values for ak37 ensembles. (E) Histograms for α torsion angle values for ak37.

To confirm this, we assessed the similarities among all MCMC and SAM ensembles of CGRP variants by using aJSD_d, a score which evaluates divergences in Cα-Cα distance distributions (**Fig 4C**). We found that the MCMC wild-type ensemble most closely resembles the wild-type one from SAM (aJSD_d=0.004) and the MCMC mutant ensemble most closely resembles the mutant one from SAM (aJSD_d=0.004). Their scores are smaller than the one between the wild-type and mutant ensembles from MCMC (aJSD_d=0.011), showing that SAM specifically models the variations. We performed a similar comparison via aJSD_t, a score to evaluate torsion angles distributions (**Fig 4D**) and observed similar trends also for these features. When we inspected the Cα-Cα distance and α angle distributions with the highest JSD between the wild-type and mutant ensembles from MCMC, we observed that SAM could correctly reproduce the differences (**S6 Fig**). Taken together, this data suggest that generative models trained on sufficiently large datasets can achieve a level of transferability necessary for accurately modeling the impact of single amino acid mutations on structural dynamics.

We also focused on ak37, the peptide for which idpSAM provides the worst approximations (**S2 Fig**). This is a synthetic peptide[45] which assumes mostly helical states in simulations. It was included here as it was part of previous simulation studies on intrinsically disordered peptide conformations[13]. Its ABSINTH ensemble is populated by fully-helical or helical hairpin conformations where two shorter helices are packed together. This results in bimodal distributions for *R*_g_ (**Fig 4F**) and Cα-Cα distances (**S7 Fig**), which idpSAM captures only approximately. Even if the model generates both types of conformations, their densities are incorrect (**Fig 4E**). IdpSAM also underestimates helicity, as seen by α angle distributions (**Fig 4G**), where values in the range of 50-60° correspond to helical backbone geometries. Our explanation is that the training set of idpSAM, which consists of naturally-occurring IDR sequences, does not contain examples with such high helicity (**S8 Fig**). The average helicity in the MCMC ensemble of ak37 is 0.70. This is an outlier in the training set, where the mean value is 0.03. However, idpSAM has enough capacity to model ak37. When we re-trained its DDPM only on ak37, the resulting model faithfully replicates its ensemble (**S9 Fig**). To model ak37 without explicitly including it in the training data, we re-trained the DDPM by adding to its full training set 5 MCMC simulations for the ak16, ak27 and ak31 peptides[45] respectively. These are shorter versions of ak37 and have ensembles with similar properties (**S3 Table** and **S10 Fig**). With this augmentation, the retrained model provides a closer approximation for ak37 (**Fig 4E** and **S7 Fig**) while maintaining good performance on the entire test set (**S4 Table**).

This analysis demonstrates how the composition of the training set determines the ability of a model to generate conformations for unseen sequences. In the case of idpSAM, it means that the sequence space is sufficiently diverse to predict the effects of mutations, but the model is limited to mostly disordered peptides and not expected to perform as well for mostly folded peptides with extensive secondary structures since such systems were not part of the training set.

### Completeness of generated conformational ensembles

The goal of protein conformational sampling is ideally to generate complete structural ensembles with correctly weighted conformations. Because higher-energy conformations do not contribute significantly to thermodynamic averages, this means in practice ensembles of all low-lying energy conformations within a few kT from the global free energy minimum. For a well-folded system, this may be a narrow set of conformations, but, for more dynamic systems such as IDRs, the relevant set of states can be quite broad. Simulations explore conformational landscapes via iterative sampling and reach new states only after crossing kinetic barriers. Therefore, short simulations almost certainly do not sample all relevant states and it may require significant effort, including enhanced sampling techniques, to generate complete converged conformational ensembles in this manner. For the training systems used here, where we used only five independent MCMC simulations for each IDR, we probably did not achieve converged sampling for many peptides.

Direct generation of conformational ensembles as with idpSAM is not subject to kinetic barriers and there is no issue in principle to generate complete, correctly weighted ensembles for all relevant states of a given system. The key question is whether this is possible with a model trained on incomplete ensembles as in this study.

To address the question of sampling completeness, we ran much more extensive sampling for the test peptides (**S2 Table**) with respect to the training ones. For a short test peptide, such as DP03125r003 (15 residues) the ensemble obtained by running five simulations is similar to the ensemble obtained from more extensive sampling from 73 simulations (**S11 Fig**). On the other hand, for longer peptides such as drk_sh3 (59 residues), ensembles from five runs can be entirely different from the extensive sampling ones (**Fig 5**) since the sampling problem becomes harder for larger molecules. In fact, for drk_sh3 we had to run replica-exchange enhanced sampling to construct the test set ensemble. Surprisingly, even though idpSAM was trained on ensembles that originated via limited sampling, it managed to approximately recover the full ensemble of drk_sh3 (**Fig 5**). The reference PCA landscape of drk_sh3, as well as its *R*_g_ and Cα-Cα distance distributions after extensive sampling are mostly captured by the model. Similar results are observed for other test peptides for which we applied enhanced sampling to reach converged ensembles (**S12 Fig**).

**Fig 5.**
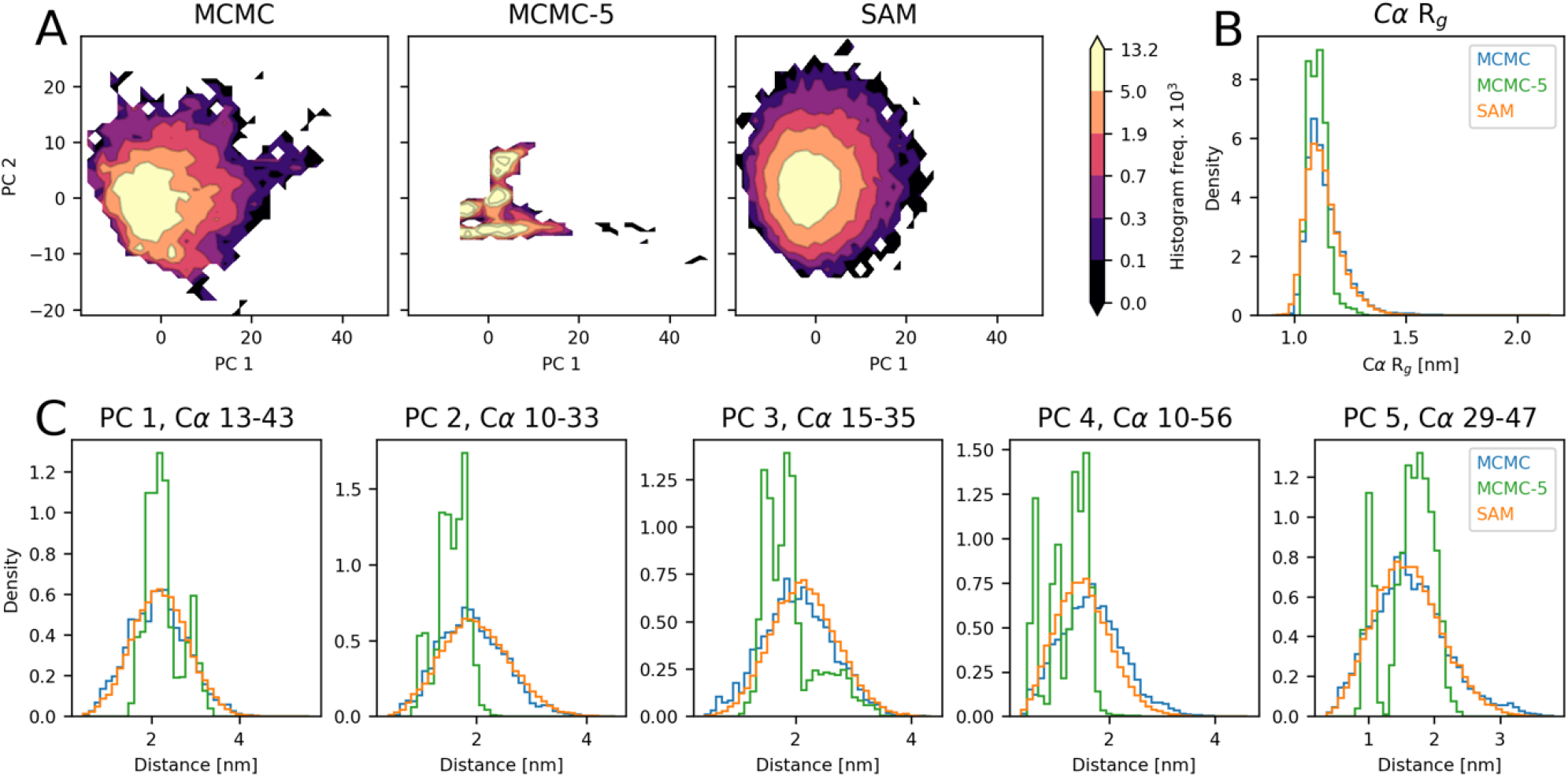
Modeling of drk_sh3 with idpSAM and different levels of MCMC sampling. (A) PCA histograms for drk_sh3 from ensembles: MCMC with extensive sampling (left), 5 MCMC runs at 298 K (MCMC-5, center) and SAM (right). Frequency values are multiplied by 1 × 10^3^. (B) Histograms of Cα *R*_g_ values for the three ensembles. (C) Histograms for selected Cα-Cα distances in the three ensembles. The distances are the ones that exhibit the highest absolute loading value for the five principal components (PCs) identified in a PCA on the MCMC ensemble from extensive sampling.

To provide a quantitative analysis, we compared the evaluation scores of idpSAM against traditional sampling via simulation, which we call the MCMC-5 strategy (**Table 1**). This strategy simply consists of generating an ensemble from 5 MCMC runs at 298 K for each test set peptide. This is the same amount of sampling that we used for the training peptides. We then compare ensembles generated by idpSAM or MCMC-5 against the converged ensembles from much more extensive sampling. For every score, idpSAM obtains better average values with respect to MCMC-5. When comparing the scores for each peptide, we find that idpSAM provides better approximations especially for longer peptides (**S13 Fig**). These results suggest that protein conformational generators such as idpSAM may efficiently use training examples with limited sampling data to provide close approximations of the full ensembles of peptides unseen during training. This means that idpSAM is not just useful for generating ensembles for different sequences but could also be used to generate more complete ensembles for systems for which limited simulation data is already available.

### Impact of training data size

We showed above that idpSAM performs better than our previous model idpGAN because of an improved generative model but also because of expanded training data. How the amount of training data translates into better performance, especially with respect to transferability, is an important question that has not yet been studied in detail for conformational generators. More specifically, it is unclear what the benefits are of including more systems vs. more sampling of the same systems, and how much additional data is necessary to improve performance by a certain amount.

To address the question of system diversity, we re-trained the DDPM of SAM with various numbers of training peptides and evaluated the models on the same test set. When training the models with different *n*_*systems*_, we kept the number of training steps constant by varying *n*_*frames*_, the number of snapshots of a system used per epoch. In this way, differences in performance are only caused by different compositions of the training sets. The general trend we observed is that performance improves by adding more training sequences. In **Fig 6A** to **E**, we show how evaluation scores improve as a function of the number of sequences. We find that modeling quality is relatively low when using less than 500 training sequences, and rapidly increases as the number of sequences reaches 1000 sequences. Adding even more sequences further improves the performance, albeit at a lower rate. This is in line with what has been observed for the OpenFold model[46]: protein modeling accuracy increases rapidly with the number of training systems and then continues to slowly improve by providing more systems.

**Fig 6.**
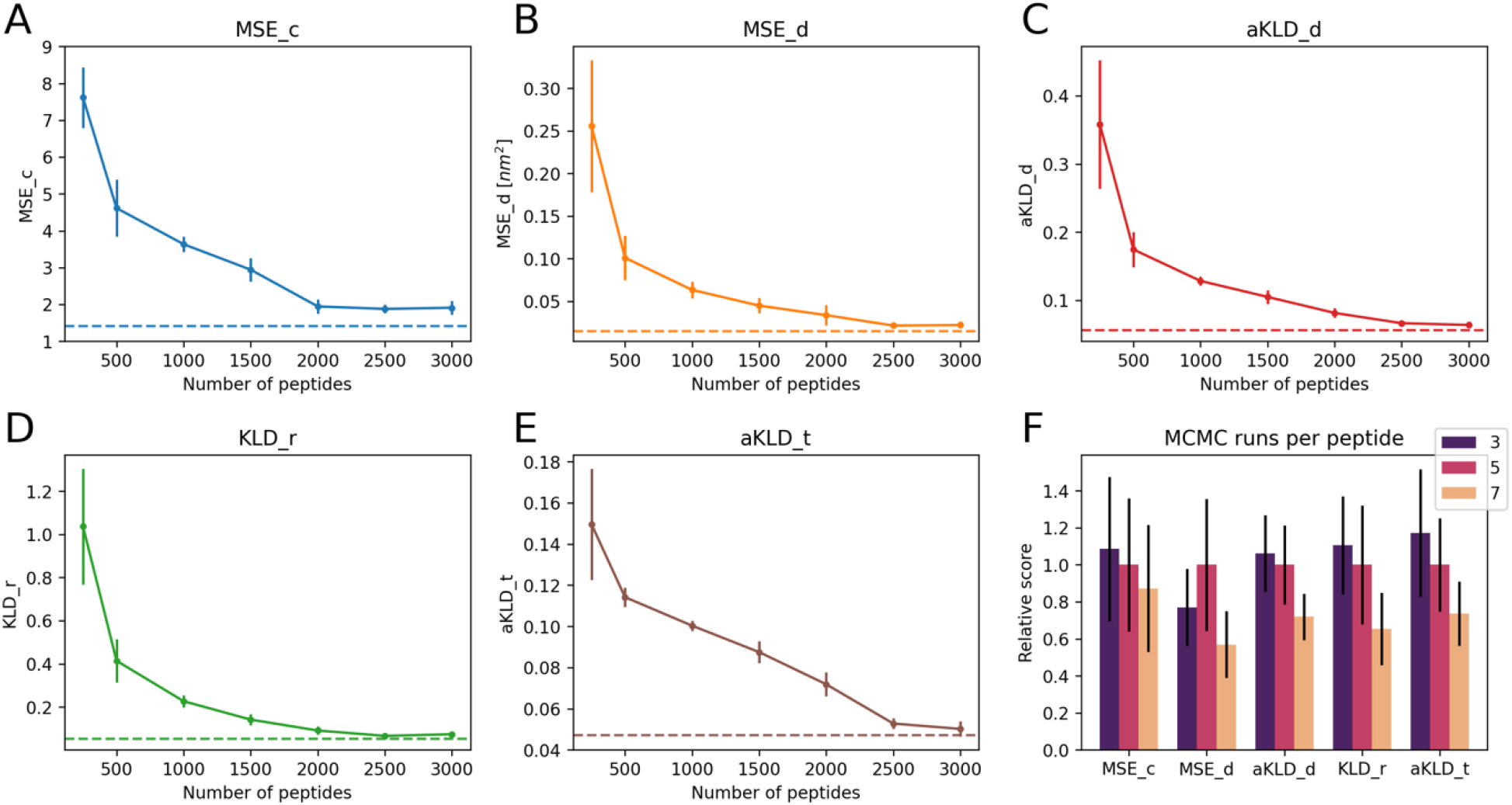
Training set dependance of SAM. (A) to (E) report the average evaluation scores for the 22 test set peptides as a function of the number of IDRs used to train SAM. Models were trained with 250, 500, 1000, 1500, 2000, 2500 and 3000 IDRs. Error bars represent the standard errors observed in 5 replicas. Each replica was trained with the same number of IDRs, but used a different random subset (uniformly sampled) from all the 3,259 training IDRs. The horizontal lines represent scores from the SAM model trained with all IDRs. Each panel shows data for a different score. (F) Comparison between SAM models trained on the set of idpGAN with 1089 IDRs. The DDPM of each model was trained with different numbers of MCMC simulations per peptide (3, 5 or 7). The heights of the bars represent the average scores obtained on the set of 22 test peptides, while error bars represent standard errors. For each score, all mean values and errors were divided by the mean value of the model trained with 5 runs per peptide, to plot different scores in similar numerical ranges.

We also investigated the effect of increasing the amount of sampling in the training set. For all the 1089 training IDRs of the ABSINTH-based idpGAN model, we performed 2 extra MCMC simulations, which we added to the 5 original ones. We trained SAM on these IDRs using 3, 5 or 7 runs, again maintaining the same number of training steps. For most evaluation scores, using more training runs appears to improve the average scores (**Fig 6F**). Given that for shorter peptides the sampling is closer to convergence, we believe that what makes a difference in this case is probably the amount of sampling for the longer peptides.

Taken together, these results illustrate the centrality of training data for protein conformational generators. Both the diversity of sequences and the amount of sampling in the training set are crucial for improving transferability. However, once a model such as idpSAM has been trained with a sufficiently large training data set, additional training data is expected to lead to diminishing returns in terms of further improved performance.

### Ablation studies and neural network capacity

Finally, we analyze the impact of different choices in model design for achieving increased performance with idpSAM. The **ϵ**_*θ*_ noise prediction network is central to the capabilities of idpSAM. To better understand the function of *ϵ*_*θ*_, we conducted ablation studies on components which we believed to be critical for its performance (**S4 Table**). The most important features are a sufficiently large number of transformer blocks and the learning rate schedule. Also the adaLN-zero mechanism for feeding conditional information[31] and the injection of the input encodings at each block have an impact, but it is less relevant.

We also tested the effect of changing the dimension *c* of residue encodings. IdpSAM uses *c* = 16. We attempted to retrain the model with *c* values of 4, 8 or 32. For values of 4 and 8, we found a decrease in modeling accuracy (**S4 Table**), which we attribute to the inferior reconstruction capability (**S1 Table**). For a value of 32, the reconstruction ability is similar to the default one. The average evaluation scores are slightly lower, though most differences do not appear to be statistically significant.

Spurred by studies showing how the performance of DDPMs scales with the expressiveness of their neural networks[47], we also explored two modifications of **ϵ**_*θ*_ (**Table 1**). First, we increased the capacity of **ϵ**_*θ*_ by changing the number of its transformer blocks from 16 to 20 (SAM-G). Then, we attempted to improve its inductive biases[48] by adding 4 FrameDiff edge update operations[26], while keeping 16 transformer blocks (SAM-E). Edge update mechanisms have been shown to be crucial for attention-based networks for proteins[4, 49]. Since they are computationally expensive, we only applied them to one out of every four transformer blocks, starting from the first. Both modifications mostly lead to slightly better average scores with respect to the default SAM, but the differences are not significant according to a Wilcoxon signed-rank test (with significance level of 0.05). Interestingly, although adding edge updates improves the validation loss (**S14 Fig**) it does not translate into significantly better ensemble modeling. The use of edge updates also comes at the cost of increased sampling times. Using the regular SAM, the average time for generating a test ensemble of 10,000 conformations is 261 s, while for SAM-E it is 1,306 s (in both cases utilizing a batch size of 256). We did not explore the possibility of combining the use of more blocks along with more edge updates, as this would have made the models excessively heavy. We believe that a key challenge in this field will be to obtain a good tradeoff between the expressiveness of models and their computational efficiency.

## DISCUSSION

The direct generation of conformational ensembles via machine learning methods is a very attractive alternative to traditional simulation approaches. The main advantage is a much greater computational efficiency when the high cost of crossing kinetic barriers via iterative sampling can be avoided. In addition, generative machine learning frameworks can be trained in principle to reproduce any target data, including experimental data, and are not limited by the specific functional forms or numerical algorithms that are commonly used in simulations. However, a key challenge is how to achieve transferability, i.e. the ability to generate meaningful ensembles for systems and/or conditions not seen in training.

Previously we focused on the generation of conformational ensembles for IDRs and showed with idpGAN that transferability to sequences unseen during training can be reached in some cases[29]. Here we demonstrate with idpSAM that consistent transferability necessary for broader applicability is possible with a diffusion model and an expanded training set. We find that idpSAM not only generates conformational ensembles for unseen sequences that closely approximate what traditional simulations would provide, but, remarkably, the generated ensembles reflect converged ensembles even though many of the training data were clearly too short to reach full convergence. This suggests that the model learned general principles about the conformational sampling of peptides that can be applied to arbitrary sequences. One way in which idpSAM could have achieved this may be by combining information acquired from shorter peptides and from incomplete sampling of longer peptides. For highly-flexible molecules like IDRs, different sequences may share at least in part their conformational space, thus making the transfer of knowledge via an effective fragment-based approach feasible.

Despite its improved transferability, our model still has some important limits, especially when modeling systems with characteristics that are uncommon in the training set. It remains an open question how to model such systems without further expanding the training set or explicitly including them in the training data. In particular, idpSAM does not give realistic ensembles for peptides that are much longer than the peptides in the training set. This is a significant limitation since many biologically relevant systems are relatively long (i.e. > 150 residues). Generating training data for very long peptides becomes computationally so demanding that it may not be possible to obtain comprehensive training data for such IDRs. In addition, increased capacity of the neural networks underlying the generative model is likely necessary to improve performance for longer peptides, which comes at increased computational costs for sampling conformations. In future research, we will explore the use of fast sampling methods[50] in conjunction with these heavier models. Another way to improve SAM could be to use probabilistic decoders. Our use of deterministic decoders is an approximation, but seems to work well empirically. We believe that this is because the encodings of SAM contain nearly complete information to reconstruct a 3D structure, therefore decoding is a one-to-one mapping problem for which a deterministic decoder is well-suited. Probabilistic decoders are instead indispensable for one-to-many mappings, like generating all-atom structures from very low-resolution CG representations[51]. The use of probabilistic decoders could also improve the general modeling capabilities of SAM. Despite these challenges, we show that transferable deep generative models are a promising strategy for modeling protein structural ensembles.

From a methodological point of view, SAM represents a novel type of latent DDPM for molecules. With SAM we observed stable training and scalability of performance with training set size. It is reasonable to think that SAM could equally work well with more complex conformational ensembles, such as those generated by explicit solvent MD simulations of IDRs or ensembles of natively more ordered peptides. We also believe a generative model like SAM could be applied also to other types of protein simulations, such as datasets of simulations of globular proteins[52]. In addition, our class of latent DDPM could be adapted to also perform all-atom modeling by using an appropriate reconstruction loss for the AE. For example, we envision the Frame Aligned Point Error[4] loss from AlphaFold2 as an attractive function for training a SAM version with atomistic details.

While idpSAM generates coarse-grained representations that can be converted to atomistic representations via other methods, such as cg2all[40], directly generating ensembles with atomistic details may ultimately give physically more realistic ensembles. This is because all-atom reconstruction methods trained on the PDB will be biased by the fact that most of the PDB data is conformationally averaged and at least in part subject to modeling that resorts to ideal stereochemical geometries when experimental data is not of sufficiently high resolution to determine otherwise. Generating atomistic ensembles also provides an avenue for including experimental data as conditional input. These directions will be the focus of future efforts.

Finally, Arts et al. demonstrated[53] that learning diffusion models is essentially very similar to learning CG force fields. Therefore, our study has also implications for the development of machine learning CG force fields for proteins, where the issue of transferability and dependance on the size of the training set are open questions[17]. Our data show that there is hope that an increase in training set size could translate into improved and more transferable neural CG force fields. It also indicates that the amount of sampling for each training system does not have to be complete in order to achieve transferability. Even as conformational generators may begin to replace traditional simulation methods, more transferable CG force fields will still be needed to address more complex biological questions, for example in systems of interacting molecules, but ultimately we see a convergence of traditional simulation approaches with machine-learning based conformational generators into new, more effective platforms for the generation of realistic conformational ensembles of complex macromolecules.

## METHODS

### Training and validation sets

The data for training idpSAM consist of ABSINTH simulations of peptides. The dataset includes data we collected previously[29] to train idpGAN. That set consisted of simulations of 1,089 peptides with lengths ranging from 20 to 50 residues. Their sequences were obtained from DisProt (version 2021_06), a manually curated database of IDRs[54]. For this study, we added simulations for another 2,170 peptides from DisProt with lengths ranging from 12 to 60. The full training set used here therefore consists of 3,259 IDRs. Details of the selection process of the sequences are reported in **S1 Text**.

The validation set is composed of ABSINTH simulations for 25 peptides (**S5 Table**) with lengths ranging from 20 to 50. We randomly extracted them from UniProtKB/Swiss-Prot[55], focusing on peptides with IDR-like properties[56] based on their sequences (**S1 Text**). We used the validation set to evaluate the loss function of the models we trained and to select the best model among training replicas (see below).

### Test set

The test set consists of ABSINTH simulations for 22 peptides (**S2 Table**). 6 of those were used before in the test set of the ABSINTH-based idpGAN model[29], while 16 are new test cases. All of the sequences belong to unstructured or synthetic peptides and their lengths range from 8 to 59 residues. None of the proteins in the test have similar sequences in the new training set. We define a query sequence to be similar to a training one if the E-value is less than 0.001 in a phmmer search[57] over the training set with default parameters. We selected the test sequences to cover different lengths and a range of net charge per residue values similar to the distribution found in training IDRs (**S8 Fig**). We used the set to evaluate the structural ensemble modeling performance of our method.

### Simulations

Simulation data was collected via Metropolis MCMC sampling of IDRs with the OPLS-AA/L force field[58] and the ABSINTH implicit solvation model[14] as implemented in the CAMPARI 4.0 package[11].

For each training and validation peptide, we performed respectively 5 and 3 independent simulations at 298 K with the same protocol that we used to collect training data for idpGAN. This protocol replicates the protocol described in the study from Mao et al.[59].

For test peptides, we employed the same protocol as for training and validation peptides, but with more extensive sampling. For each test peptide we ran as many simulations as needed to reach converged sampling (**S2 Table**). The rationale was to evaluate the ability of the generative model to recover the converged distributions of the test peptides even though training data may not be fully converged. For four of the longest peptides (nls, protan, protac and drk_sh3), we additionally performed enhanced sampling via thermal replica exchange (RE) MCMC[30, 60], since we could not reach convergence with regular constant temperature simulations. Details of all CAMPARI protocols in this study are reported in **S2 Text**.

### Generative model framework

IdpSAM makes use of a DDPM as a generative model. DDPMs attempt to approximate probability distributions by learning the reverse of a diffusion process over the data. We adopted the probabilistic and parameterization framework[19] introduced by Ho et al. 2020, which we refer to for a detailed treatment of DDPMs. Briefly, the goal of DDPM training is to learn to sample from an unknown distribution *q*(**x**_0_) describing the elements **x**_0_ in the training data. This is commonly achieved by training a noise prediction network ϵ_*θ*_ to denoise training set objects perturbed with various levels of random Gaussian noise. Once the network is trained, it can be used to generate samples distributed approximately as *q*(**x**_0_) by gradually denoising samples extracted from a normal distribution *N* (**0, I**). In this work, we are interested in modeling the distribution of the Cartesian coordinates of Cα atoms of peptides. The latent DDPM strategy that we explore here proceeds as follows: (i) we first train an autoencoder (AE) to learn an Euclidean-invariant encoding of the coordinates of Cα atoms of proteins; (ii) we then train a DDPM on encoded Cα coordinates; (iii) finally, for generating samples at inference time, we sample from the DDPM and map the generated encodings to the corresponding Cα coordinates using the decoder part of the AE.

### Invariant structural AE

The first stage of training our latent DDPM is to train an AE. The goal is to learn to encode Cα traces from protein structures into a representation that can be easy to model with a DDPM. We define an encoder function *E*_*ϕ*_ with learnable parameters *ϕ* as:

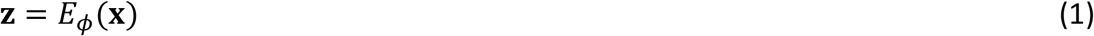

where *x* ∈ ℝ^*L×*3^ are the Cartesian coordinates of the Cα atoms of a protein with *L* residues and *z* ∈ ℝ^*L×c*^ is an encoding (that is, a “latent” representation) for that conformation. Importantly, to simplify the learning task of the downstream DDPM, we aim to learn SE(3)-invariant encodings. Formally:

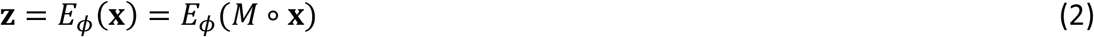

requiring that any rigid transformation *M* consisting of rotations and translations in Euclidean space preserves the object and its orientation[4, 26] (i.e., a distortion or reflection of the coordinates is not permitted). Therefore, these encodings can be viewed as a form of learnable internal coordinates. This is different from other latent DDPM implementations for molecules[34, 35], where the encodings are points in Euclidean space.

The other component of the AE is a decoder function *D*_*ψ*_ with learnable parameters *ψ*:

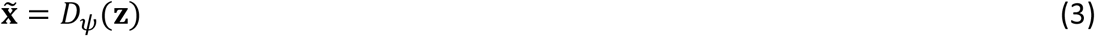

where 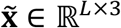is a reconstructed conformation. To train an AE, the objective function is based solely on reconstruction:

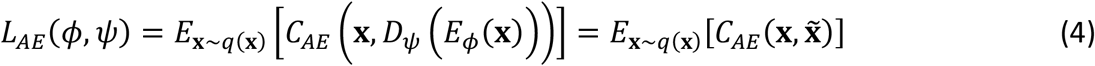

where *q* (**x**) is the distribution of conformations in the training set. The reconstruction loss for a single conformation is composed of two terms:

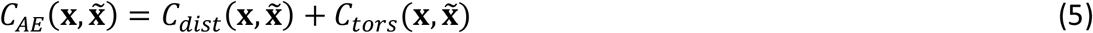

*C*_*dist*_is a term for evaluating the reconstruction of all Cα-Cα distances in the molecule. *C*_*tors*_ is a reconstruction term for the torsion angles formed by four consecutive Cα atoms in the molecule, which we term as the α angles. The first term is a mean squared error (MSE) defined as:

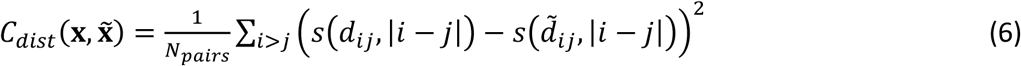

where *N*_*pairs*_ = *L* (*L −* 1)/2 is the number of Cα-Cα distances in a peptide, *d*_*ij*_ = *‖x*_*i*_ *– x*_*j*_*‖* and 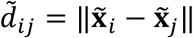 are the original and reconstructed Euclidean distances between Cα atoms *i* and *j* and *s* is a standard scaler linear function with the sole purpose of numerically helping with training (**S3 Text**). Because the set of all pairwise distances in a 3D point cloud can be used to define the position of the points up to a E(3) transformation[61, 62], the *C*_*dist*_term is in principle sufficient to obtain perfect molecular reconstructions up to a reflection. However, proteins are chiral molecules[63]. To break the reflection symmetry in the learned representations, one possibility would be to rely on SE(3)-invariant neural networks[64, 65]. Instead, we adopted a simple approximation which consists in forcing the AE to learn the correct mirror images by using a reconstruction loss for torsion angles α, which are molecular features with a chiral distribution[29, 66]. The term is expressed as:

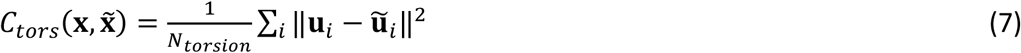

where *N*_*torsion*_ = (*L −* 3) is the number of torsion angles in a peptide chain, ***u***_*i*_ and 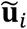 are vectors[4] storing the cosine and sine values of the torsion angle between Cα atoms *i, i +* 1, *i +* 2 and *i +* 3 of training and reconstructed conformations, respectively. Empirically, we found that AEs trained with *L*_*AE*_ have high reconstruction accuracy and learn to correct mirror images (see Results). Therefore, *E*_*ϕ*_ can be viewed as an SE(3)-invariant encoder for practical purposes. Finally, we note that the reconstruction objective considers only invariant geometrical features. We do not aim to recover the rigid-body translation and rotation of the original conformation. This choice allows us to circumvent the use of an additional equivariant function to store reconstruction group actions, as prescribed by Winter at al. for geometrical autoencoders[34, 67].

### AE neural networks

The use of invariant encodings gives freedom over the choice of the architectures for *E*_*ϕ*_ and *D*_*ψ*_. For both networks, we use transformer architectures[36], which work well for protein data[68] and naturally allow modeling protein chains with different lengths.

The *E*_*ϕ*_ network takes as input the full distance matrix and the α torsion angles of a conformation **x**. These features are all internal coordinates, which is what allows encodings to be translationally and rotationally invariant[69]. The output of *E*_*ϕ*_ is an encoding **z**. We set *c*, the dimension of a Cα encoding, to 16. At first sight, it may seem counterintuitive to represent Cα atoms via vectors with more dimensions than the 3 original Cartesian coordinates. However, our goal is not to learn an efficient Euclidean-invariant dimensionality reduction for protein structures, rather to optimize the performance of a latent DDPM. Empirically, we found *c* = 16 to work best for our data (see Results).

As for *D*_*ψ*_, the network takes as input an encoding and maps it back to 3D coordinates. Full details of the two networks are reported in **S3 Text** and **S15 Fig**.

### Latent DDPM

Once an AE is trained, it is possible to use *E*_*ϕ*_ to encode training conformations and to apply a DDPM to learn their distribution. Here, the DDPM is a conditional model. Its goal is to learn *q* (**z**_0_|**a**), that is, the distributions of encoded conformations **z**_0_ for a peptide with amino acid sequence **a**, which represents the condition. We train our model with the variational lower bound objective *L*_*simple*_ commonly used in DDPM applications[19]:

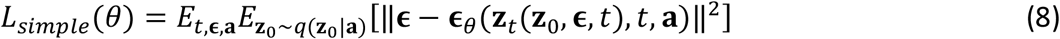

Here 1 *< t ≤ T* is an integer-valued diffusion time step and is extracted from a uniform distribution. *T* is the total number of steps of the diffusion process. **ϵ** ∈ ℝ^*L×c*^ is random noise extracted from *N* (**0, I**) used to perturb an encoded training set conformation **z**_0_. **a** is an amino acid sequence associated with a conformation and is extracted from the distribution of sequences in the training set. **z**_*t*_ is the perturbed version of **z**_0_, which can be computed in closed form[19] as a function of **z**_0_, **ϵ**, and *t*. The task of the **ϵ**_*θ*_ network is, given its input *z*_*t*_, to predict the **ϵ** which can be used to reconstruct *z*_0_.

### DDPM neural network

The noise prediction network **ϵ**_*θ*_ takes as input a perturbed encoding **z**_*t*_, the corresponding perturbation step *t* and the amino acid sequence **a** of the peptide. The architecture of **ϵ**_*θ*_ is again a transformer. We incorporated in the network several modifications inspired by recent works where transformers are used in latent DDPMs. The most important are the use of the adaLN-Zero mechanism[31] for injecting timestep and amino acid embeddings in the network and also the injection of the input perturbed encoding at each transformer block. Full details for **ϵ**_*θ*_ are reported in **S3 Text** and **S16 Fig**.

### Sampling and decoding

For the diffusion process during training, we use *T* = 1,000. We adopt a sigmoid noise schedule[70] for the process. When sampling at inference time, we use 100 steps via the accelerated generation algorithm introduced with denoising diffusion implicit models[42]. Unless otherwise stated, all results presented in the article are obtained by generating samples with 100 steps.

For decoding the generated encoded conformations, we directly use the *D*_*ψ*_ network obtained in the AE training. In our work, the decoder is therefore a deterministic function. In theory, to perfectly replicate the distribution of objects **x**_0_ (in our case, Cα coordinates), the decoder of a latent generative model should be probabilistic[33], as for example variational autoencoders[21]. However, we discovered that this approximation to work well in practice and discuss this choice in the Results section.

### Training process

For training the AE and the DDPM we define two parameters called *n*_*systems*_ and *n*_*frames*_. The first is the total number of peptides used in training. With our full training set *n*_*systems*_ = 3,259. The second parameter is the number of snapshots randomly extracted from the MCMC data of each system during a training epoch. We use different *n*_*systems*_ and *n*_*frames*_ values for the AE and the DDPM (the former is trained with only a part of the full dataset to speed up training) and different number of epochs. When training, we always adopt batches containing protein conformations with the same number of residues to avoid padding strategies^23^.

During AE training, we perturb input conformations with Gaussian noise with *σ* = 0.1 Å as a form of data augmentation which we found to help the learning process.

Both models are trained with the Adam optimizer. We found the use of learning rate schedules to be critical in training and used different schedules for each model (see Results section).

When training a model, we run multiple replicas and, at the end of training, we select the one with the best loss over the validation set. Full details of the training processes and their computational times are reported in **S6 Table**.

### Implementation

All neural networks and deep learning algorithms in this work are implemented using the PyTorch library[71]. We employ the Diffusers library[72] to run the forward (noising) and reverse (generative) diffusion processes. The code and weights for a pre-trained idpSAM are available at https://github.com/giacomo-janson/idpsam.

### Evaluation

To compare the properties of ensembles generated by SAM (or other strategies) with the reference MCMC ones, we used different scores based on previous work[29]. Here we provide brief descriptions. More details are given in **S4 Text**. The MSE_c score evaluates the similarity between reference and generated Cα-Cα contact map frequencies, with 8 Å as a contact threshold. MSE_d compares average Cα-Cα distance values. aKLD_d evaluates an approximation of the Kullback-Leibler divergence (KLD) between sets of reference and generated Cα-Cα distance distributions. aKLD_t similarly evaluates the KLDs between α angle distributions. The aJSD_d and aJSD_t are similar to the last two scores, but instead use the symmetric Jensen-Shannon divergence (JSD). We use them to compare two ensembles when we do not consider one of them to be a reference. Finally, KLD_r evaluates the distribution of Cα radius-of-gyration (*R*_*g*_) values.

For each test set peptide, we generated with SAM an ensemble of 10,000 conformations and compared it via these scores against an ensemble composed of the same number of snapshots from MCMC data. In case of the test peptides for which we did not employ RE, MCMC conformations were randomly extracted from all the available simulations at 298 K. For those peptides with RE data, 25% of the snapshots were extracted from constant temperature simulations at 298 K, the remaining 75% from the replicas at 298 K in the RE simulations.

All plots in the manuscript were generated for ensembles with 25,000 conformations obtained using this protocol from MCMC data or generated by SAM. To visualize the distribution of conformations, we constructed histograms of features identified in principal component analysis (PCA). We performed PCA on the reference MCMC ensembles using all Cα-Cα distances as input features and then projected conformations from the generated ensembles onto the principal components obtained from the reference simulation data.

The 3D structures in this work were analyzed and rendered via NGLview[73].

## Data Availability

The neural network code and weights for a pre-trained idpSAM as well as training, test, and validation sequences are available at https://github.com/giacomo-janson/idpsam [74].

## Acknowledgements

Funding from NIH R35 GM126948 is acknowledged. This work used Expanse (at San Diego Supercomputer Center), Bridges-2 (at Pittsburgh Supercomputing Center) and Rockfish (at Johns Hopkins University) resources through allocation BIO230084 from the Advanced Cyberinfrastructure Coordination Ecosystem: Services & Support (ACCESS) program[75], which is supported by National Science Foundation grants #2138259, #2138286, #2138307, #2137603, and #2138296. High-performance computing resources were also used at the Institute for Cyber Enabled Research (ICER) at Michigan State University.

The authors thank Andreas Vitalis for his kind help with the CAMPARI package and the ABSINTH model and thank Alexander Dickson and Ceren Kılınç for having provided the idea to model CGRP sequence variants.

