## Supplementary Information for "Transferable deep generative modeling of intrinsically disordered protein conformations"

### Supplementary Text

#### S1 Text. Selection of sequences.

##### Selection of the training set sequences

The training set in this study consists of 3,259 IDR sequences. The set was constructed via an incremental process involving the addition of different parts, which are described here.

1. **1089 IDRs from the training set of ABSINTH-based idpGAN.** The first part of the training set consists of data we previously collected to train the idpGAN model[1]. Two different idpGAN models were trained on either COCOMO coarse-grained simulations[2] or ABSINTH implicit solvent simulations[3]. The COCOMO-based idpGAN was trained on a set of 1,966 IDRs. These sequences originate from DisProt[4] (version 2021_06) and have lengths ranging from 20 to 200 residues. The ABSINTH-based idpGAN model was trained on a subset of these, containing all the 1089 sequences with lengths between 20 and 50 residues. This subset, for which we collected ABSINTH simulation data, is the initial part of the training set of the present study.
2. **132 IDR sequences from the training set of COCOMO-based idpGAN.** In addition to the sequences of part 1, we selected all 132 IDRs with lengths from 51 to 60 from the training set of COCOMO-based idpGAN. The goal here was to increment the maximum length of the peptides in the training set to obtain generative models able to work with longer peptides.
3. **1,888 IDR random crops from the training set of COCOMO-based idpGAN.** To further expand the training set, we added 1,888 IDR sequences with lengths ranging from 20 to 55. These sequences were generated by randomly extracting one or more continuous crops from of each IDR in the training set of COCOMO-based idpGAN with a length between 56 and 200. The length of the crops was randomly sampled from a uniform distribution. In the extraction process, we ensured that no two crops taken from the same original IDR had overlapping sequences. The goal was to significantly expand the diversity of sequences with intermediary lengths in the training set.
4. **150 IDR sequence with lengths between 12 and 19.** Finally, we randomly selected from DisProt (see part 1) a new set of 150 IDRs with 12 to 19 residues. The goal was to include a portion of shorter sequences in the training set for improving modeling performance on shorter peptides.

We ran ABSINTH MCMC simulations for all IDR sequences of part 2 to 4 using the same protocol used for part 1. The resulting data was added to the data from part 1 to finally constitute the full training set of this study.

##### Selection of the validation set sequences

The validation set in this study consists of 25 peptides. To obtain their sequences, we used the Swiss-Prot database[5] containing a total of 568,744 protein sequences (retrieved on February 2023). For each sequence with at least 20 amino acids, we extracted a random crop with length between 20 and 55 (uniformly sampled). We then filtered the sequences via simple and approximate criteria for IDP classification based on charge-hydropathy values[6]. We kept all crops classified a disordered and a random 20% fraction of the crops classified as not disordered. From all this sequences, we randomly selected 25 for constituting the validation set. For these peptides, we ran MCMC simulations using the same protocol employed for the training set.

#### S2 Text. MCMC simulation protocol.

Here we describe the protocols used to run MCMC simulations via CAMPARI[7] in this study, which was adopted from a previous study for simulating charged IDRs[8].

##### MCMC runs at 298 K

**Hamiltonian**: simulations were performed using the OPLS-AA/L force field and the ABSINTH implicit solvent model. The CAMPARI parameter file *abs3.1_opls.prm* was used. Cutoffs for Lennard-Jones and electrostatic interactions were set at 10 and 14 Å respectively.
**Peptide modeling**: peptides were capped with an acetyl group at the N-terminus and a N-methylamide group at the C-terminus. Histidine sidechains were protonated only at the ε position, making them neutrally charged.
**System**: peptides were placed inside a spherical droplet. The radius of the droplet was set to 70 Å for peptides with $L$ <= 34 residues ("short" peptides) and 100 Å for peptides with $L$ >= 35 residues ("long" peptides). $Na^+$ and $Cl^-$ ions were added to neutralize net peptide charges and to represent a 125 mM salt solution (resulting in 108 excess ion pairs for 70 Å droplets and 315 pairs for 100 Å droplets).
**Sampling**: Metropolis MCMC simulations were performed in the NVT ensemble at 298 K. The degrees of freedom in the simulations were backbone ϕ, ψ, ω torsion angles and sidechain χ torsion angles and rigid-body coordinates for peptides molecules and ions.
**Monte Carlo move set**: the move set was based on a one employed previously[8].
**Simulations**: independent simulations were performed using randomly generated initial conformations. For "short" peptides, we performed $1 \times 10^6$ equilibration and $2.5 \times 10^7$ production steps. For "long" peptides, we performed $2 \times 10^6$ equilibration and $5 \times 10^7$ production steps.
**Output**: snapshots were saved every 5,000 steps during the production phase, resulting in 5,000 and 10,000 snapshots for a "short" and "long" peptide simulation respectively.

##### Replica exchange (RE) simulations

For four test set peptides (nls, protan, protac and drk_sh3), we additionally performed thermal RE MCMC simulations as implemented in CAMPARI. The RE strategy was inspired by one used previously to simulate polyampholytic peptides[9]. The temperature schedule comprised at least 12 temperatures: [298K, 302K, 306K, 310K, 315K, 320K, 330K, 340K, 350K, 360K, 370K, 380K]. For all simulations of nls and drk_sh3 and a small portion of protan and protac simulations, we added 4 temperatures: [390K ,400K, 410K, 420K]. The simulation setup (Hamiltonian, system preparation and MCMC sampling) was the same described above for simulations at constant 298 K. Each thermal replica was initiated using a randomly generated initial conformations. For most RE simulations, we performed $5 \times 10^5$ equilibration and $1.25 \times 10^7$ production steps. In a minority of cases, we performed $1 \times 10^6$ equilibration and $2.5 \times 10^7$ production steps. Swaps between two neighboring replicas were always attempted every 2,000 steps. Snapshots were saved every 5,000 steps.

### S3 Text. Neural networks of idpSAM.

This section describes the neural networks of idpSAM with pseudocode for some key operations. The PyTorch code for all networks can be found at: https://github.com/giacomo-janson/idpsam.

#### Encoder

The encoder network $E_{\phi}$ of is based on a transformer architecture (**S15 Fig A** and **B**). It has 912,888 trainable parameters.

**Input**. The input of $E_{\phi}$ consist of: (i) two types of geometrical features extracted from the coordinates $\mathbf{x} \in \mathbb{R}^{L \times 3}$ of the Cα atoms of a peptide ($L$ is the number of its residues); (ii) the amino acid sequence tokens of the peptide $\mathbf{a} \in \mathbb{R}^{L \times 1}$. The first type of geometrical features consists in the full distance matrix $\mathbf{d} \in \mathbb{R}^{L \times L}$ calculated from $\mathbf{x}$. This is processed by a radial base function (RBF) expansion as in SchNet[10, 11] with 320 equally spaced Gaussians from 0.0 to 30.0 Å. The RBF expansion is embedded to a dimension of $c_{dist} = 192$ by a multilayer perceptron (MLP) with a GELU activation[12]:

$$\mathbf{h}_{dist} = \mathrm{Linear}(\mathrm{GELU}(\mathrm{Linear}(\mathrm{RBF}(\mathbf{d})))) \qquad \mathbf{h}_{dist} \in \mathbb{R}^{L \times L \times c_{dist}}$$

The second type of geometrical features is a sequence $\mathbf{t} \in \mathbb{R}^{L \times 3}$ storing α torsion angle values (see main text). The first two channels of $\mathbf{t}$ store the cosine and sine values of the $L - 3$ angles of a peptide, with the first and last two positions padded with 0. The third channel contains a mask with values of 0 for the three zero-padded positions and 1 for the rest. These features are embedded to a dimension of $c_{node} = 128$ by an MLP:

$$\mathbf{h}_{tors} = \mathrm{Linear}(\mathrm{GELU}(\mathrm{Linear}(\mathbf{t}))) \qquad \mathbf{h}_{tors} \in \mathbb{R}^{L \times c_{node}}$$

The amino acid sequence $\mathbf{a}$ is processed via a learnable embedding layer[1, 13] to yield an amino acid embedding $\mathbf{h}_{aa} \in \mathbb{R}^{L \times c_{aa}}$ (with $c_{aa} = 32$).

**Transformer blocks**. The network has $n_{blocks} = 5$ transformer blocks[14] with self-attention and $n_{heads} = 8$ heads, a “pre” layer normalization configuration[15], an MLP with hidden dimension of 256, GELU activation and no dropout. The input of the first block is $\mathbf{h}^{[1]} = \mathbf{h}_{tors}$. At each block $l$, a linear layer prepares the input of the block in the following way:

$$\mathbf{h}_{in}^{[l]} = \mathrm{Linear}(\mathrm{concat}(\mathbf{h}^{[l]}, \mathbf{h}_{aa})) \qquad \mathbf{h}^{[l]} \in \mathbb{R}^{L \times c_{node}}, \mathbf{h}_{in}^{[l]} \in \mathbb{R}^{L \times c_{node}}$$

where $\mathbf{h}^{[l]}$ is the input sequence at block $l$.
Also, at every block the distance features $\mathbf{h}_{dist}$ are concatenated to a learnable 2d relative positional embedding $\mathbf{h}_{pos2d} \in \mathbb{R}^{L \times L \times c_{pos2d}}$ (with $c_{pos2d} = 64$) used in AlphaFold2 and other models[1, 13]. These concatenated features are projected via a linear layer to obtain:

$$\mathbf{h}_{bias}^{[l]} = \mathrm{Linear}(\mathrm{concat}(\mathbf{h}_{dist}, \mathbf{h}_{pos2d})) \qquad \mathbf{h}_{bias}^{[l]} \in \mathbb{R}^{L \times L \times n_{heads}}$$

which contains bias terms that are then added to the logit values of the attention heads in the layer. In summary, the transformer block operates the following update:

$$\mathbf{h}^{[l+1]} = \mathrm{Transformer}(input = \mathbf{h}_{in}^{[l]}, bias = \mathbf{h}_{bias}^{[l]})$$

**Output**. The output of the last layer if processed via:

$$\mathbf{z} = \text{LayerNorm}\big(\text{Linear}(\text{GELU}(\text{Linear}(\mathbf{h}^{[n_{blocks}]})), out\_dim = c)\big) \qquad \mathbf{z} \in \mathbb{R}^{L\times c}$$

where $z$ is the encoded representation of the Cα coordinates of the peptide and $c = 16$. The layer normalization operation does not use learnable elementwise affine parameters. It is used to rescale the output of the encoder to help downstream applications with other neural networks.

#### Scaling of interatomic distances in the AE loss

The AE loss used in this study includes a term for the reconstruction of Cα-Cα interatomic distances. Before calculating the loss, a distance $d_{ij} = \|\mathbf{x}_i - \mathbf{x}_j\|$ between the Cα atoms of residue $i$ and $j$ with sequence separation $k = |i - j|$ is first scaled by:

$$s(d_{ij}, k) = \frac{(d_{ij} - m_k)}{s_k}$$

where $m_k$ and $s_k$ are the mean and standard deviation in the AE training set for Cα-Cα distances with sequence separation $k$. We found this standardization procedure to be critical for efficiently training the AE model, since the scale of distances between atom pairs with different $k$ values can vary greatly.

#### Decoder

The decoder $D_\psi$ of SAM is similar to the encoder (**S15 Fig C**). It has 1,161,347 trainable parameters. In describing its components, we employ similar notations to those used for the encoder, for reasons of economy and clarity.

**Input**. The input of the decoder is an encoding $\mathbf{z}$. This is first projected to a dimension of $c_{node} = 128$ by an MLP:

$$\mathbf{h}_{enc} = \text{Linear}(\text{GELU}(\text{Linear}(\mathbf{z}))) \qquad \mathbf{h}_{enc} \in \mathbb{R}^{L\times c_{node}}$$

which is the input to the first transformer block, so $\mathbf{h}^{[1]} = \mathbf{h}_{enc}$.

**Transformer blocks**. The network has a stack of $n_{blocks} = 5$ modified transformer blocks. They share similar architecture and hyper-parameters of the blocks in $E_\phi$, with only one exception: instead of using a scaled dot product attention[14] with 8 heads, they use $n_{heads} = 32$ heads with an attention mechanism inspired by the kernel self-attention in the Timewarp model[16] and AF2 invariant point attention[13]. More specifically, the input $\mathbf{h}^{[l]}$ of a block $l$ is first mapped to query and key 3D coordinates:

$$\mathbf{q}^{(m)} = \text{Linear}(\mathbf{h}^{[l]}) \qquad \mathbf{q}^{(m)} \in \mathbb{R}^{L\times 3}$$
$$\mathbf{k}^{(m)} = \text{Linear}(\mathbf{h}^{[l]}) \qquad \mathbf{k}^{(m)} \in \mathbb{R}^{L\times 3}$$

where $m$ is the index of the head (we omit the block index in the notations for $\mathbf{q}$ and $\mathbf{k}$). Value vectors are instead obtained with the usual mechanism used in dot product attention. The logit values $y_{ij}^{(m)}$ of the attention map are calculated by:

$$y_{ij}^{(m)} = -\frac{\|\mathbf{q}_i^{(m)} - \mathbf{k}_j^{(m)}\|^2}{l^{(m)}} \qquad y_{ij}^{(m)} \in \mathbb{R}$$

with:

$$l^{(m)} = \text{softplus}(\lambda^{(m)} + \epsilon)$$

where $\lambda^{(m)}$ is a learnable scaling parameter and $\epsilon$ is a small real number for numerical stability. Similar to the encoder, the decoder uses a layer for creating 2d relative positional embeddings with $c_{pos2d} = 64$. These are linearly projected to obtain bias values $\mathbf{h}_{bias}^{[l]} \in \mathbb{R}^{L\times L\times n_{heads}}$ which are summed to logits to obtain final attention maps. Like in dot product attention, the maps are then used to update $\mathbf{h}^{[l]}$ by matrix multiplication with the values vectors. Overall, the update is:

$$\mathbf{h}^{[l+1]} = \text{ModifiedTransformer}(input = \mathbf{h}^{[l]}, bias = \mathbf{h}_{bias}^{[l]})$$

We found the use of this self-attention mechanism to slightly improve the reconstruction accuracy of the decoder. We hypothesize that the improvement could be caused by better inductive biases for modeling Cartesian coordinates.

**Output**. The output of the final transformer block is ultimately mapped to a tensor $\tilde{\mathbf{x}} \in \mathbb{R}^{L\times 3}$ by an MLP:

$$\tilde{\mathbf{x}} = \text{Linear}(\text{GELU}(\text{Linear}(\mathbf{h}^{[n_{blocks}]})), out\_dim = 3)$$

which represent the reconstructed Cα coordinates of an encoding $\mathbf{z}$.

#### Noise prediction network

The noise prediction network $\boldsymbol{\epsilon}_\theta$ of SAM is also based on a transformer architecture (**S16 Fig**). It has 15,161,296 trainable parameters. In describing its components, we again employ similar notations to those used for the other networks.

**Input**. The input of $\boldsymbol{\epsilon}_\theta$ consist of: (i) an encoded peptide conformation $\mathbf{z}_t \in \mathbb{R}^{L\times c}$ perturbed at an integer-valued timestep $1 < t \leq T$; (ii) the timestep $t$; (iii) the amino acid sequence $\mathbf{a}$ of the peptide. The input encoding is first project to the node hidden dimension $c_{node} = 256$ by a linear layer:

$$\mathbf{h}_{enc} = \text{Linear}(\mathbf{z}_t) \qquad \mathbf{h}_{enc} \in \mathbb{R}^{L\times c_{node}}$$

which is the input to the first transformer block, so $\mathbf{h}^{[1]} = \mathbf{h}_{enc}$. The input timestep $t$ is processed via a sinusoidal embeddings and an MLP with a SiLU activation to project it to a dimension of $c_{time} = 256$, then is finally tiled to a sequence of $L$ tokens:

$$\mathbf{h}_{time} = \text{Linear}(\text{SiLU}(\text{Linear}(\text{SinusoidalEmbed}(t, dim = 256)))) \qquad \mathbf{h}_{time} \in \mathbb{R}^{1\times c_{time}}$$
$$\mathbf{h}_{time} \leftarrow \text{tile}(h_{time}, dim = 0) \qquad \mathbf{h}_{time} \in \mathbb{R}^{L\times c_{time}}$$

Similar to the encoder, $\boldsymbol{\epsilon}_\theta$ uses a layer for creating amino acid embedding $\mathbf{h}_{aa} \in \mathbb{R}^{L\times c_{aa}}$ (with $c_{aa} = 32$).

**Transformer blocks**. The network has $n_{blocks} = 16$ modified transformer blocks with self-attention with $n_{heads} = 16$ heads, a "pre" layer normalization configuration, an MLP with hidden dimension of 512, GELU activation and no dropout. The modification consists in the way timestep and amino acid embeddings are injected. We use the adaLN-Zero (LN: layer normalization) mechanism from the

Latent Diffusion Transformer[17]. The scale ($\boldsymbol{\gamma}$), shift ($\boldsymbol{\beta}$) and gate ($\boldsymbol{\alpha}$) values for adaLN-Zero are obtained by:

$$\mathbf{c}_{in} = \mathbf{h}_{time} + \mathrm{Linear}(\mathbf{h}_{aa}) \qquad \mathbf{c}_{in} \in \mathbb{R}^{L\times c_{time}}$$
$$\boldsymbol{\gamma}_1, \boldsymbol{\beta}_1, \boldsymbol{\alpha}_1, \boldsymbol{\gamma}_2, \boldsymbol{\beta}_2, \boldsymbol{\alpha}_2 = \mathrm{Linear}\big(\mathrm{GELU}(\mathbf{c}_{in})\big) \qquad \boldsymbol{\gamma}_1, \boldsymbol{\beta}_1, \boldsymbol{\alpha}_1, \boldsymbol{\gamma}_2, \boldsymbol{\beta}_2, \boldsymbol{\alpha}_2 \in \mathbb{R}^{L\times c_{node}}$$
$$\boldsymbol{\zeta}_{in} = (\boldsymbol{\gamma}_1, \boldsymbol{\beta}_1, \boldsymbol{\alpha}_1, \boldsymbol{\gamma}_2, \boldsymbol{\beta}_2, \boldsymbol{\alpha}_2)$$

these values are used to normalize the embedded sequence in the transformer block (we omit the block index in their notations). We found the use of adaLN-Zero mechanism highly beneficial for the performance of SAM (see the main text), which is consistent with Peebles and Xie[17]. The injection mechanism of the adaLN-Zero we implement is illustrated in **S16 Fig B** and the gate and modulate algorithms are given below (the $\odot$ operator indicates element-wise multiplication and addition with 1 involves broadcasting):

$$\textbf{def}\ \mathrm{modulate}\big(\mathbf{h} \in \mathbb{R}^{L\times c_{node}}, \boldsymbol{\gamma} \in \mathbb{R}^{L\times c_{node}}, \boldsymbol{\beta} \in \mathbb{R}^{L\times c_{node}}\big):$$
$$\qquad \textbf{return}\ \mathbf{h} \odot (1+\boldsymbol{\gamma}) + \boldsymbol{\beta}$$

$$\textbf{def}\ \mathrm{gate}\big(\mathbf{h} \in \mathbb{R}^{L\times c_{node}}, \boldsymbol{\alpha} \in \mathbb{R}^{L\times c_{node}}\big):$$
$$\qquad \textbf{return}\ \mathbf{h} \odot \boldsymbol{\alpha}$$

Note that the layer normalization operations preceding the adaLN-Zero operations do not use learnable element wise affine parameters. Similar to both AE networks, $\boldsymbol{\varepsilon}_\theta$ also uses a layer for creating 2D relative positional embeddings with $c_{pos2d} = 64$. Again, these are linearly projected to bias values $\mathbf{h}_{bias}^{[l]} \in \mathbb{R}^{L\times L\times n_{heads}}$ which are summed to logits to obtain attention maps for a regular multi-head self-attention mechanism. The modified transformer block update can be summarized as:

$$\mathbf{h}_{out}^{[l]} = \mathrm{LatentDiffusionTransformer}(input = \mathbf{h}^{[l]}, bias = \mathbf{h}_{bias}^{[l]}, adalnzero\_input = \boldsymbol{\zeta}_{in}^{[l]})$$

to complete the update, the input of the first transformer block is used to carry out the following operation at every block:

$$\mathbf{h}^{[l+1]} = \mathrm{LayerNorm}(\mathbf{h}_{out}^{[l]} + \mathrm{Linear}(\mathbf{h}^{[1]}))$$

We found the injection of the input embedding at every block to be beneficial for SAM performance.

**Output**. The output of the last block if processed via:

$$\hat{\boldsymbol{\epsilon}} = \mathrm{Linear}(\mathrm{GELU}(\mathrm{Linear}(\mathbf{h}^{[n_{blocks}]})), out\_dim = c) \qquad \hat{\boldsymbol{\epsilon}} \in \mathbb{R}^{L\times c}$$

where $\hat{\boldsymbol{\epsilon}}$ is the predicted noise.

##### Standardization of encodings

To numerically help DDPM training, each of the $c$ channels in a training set encoding is normalized via a standard scaler using the mean and standard deviation of the channel in the dataset. At inference time, the encodings generated by the DDPM are first transformed back by the inverse of the standard scalar function. Then they are used as input for the decoder. Note that the encodings produced by $E_\phi$ are already normalized via a layer normalization operation in the output module (see above), but we found this additional standardization procedure to slightly improve the training stability of the DDPM.

#### S4 Text. Evaluation metrics.

For comparing a proposed structural ensemble (e.g.: from SAM) of a peptide with a reference MCMC ensemble of the same peptide, we used different evaluation scores. In this work, they are calculated using ensembles with 10,000 conformations each.

**MSE_c**

The Cα-Cα contact probabilities for a peptide of length $L$ were evaluated with the following mean squared error (MSE) score:

$$\text{MSE\_c} = \frac{1}{N_{pairs}} \sum_{j-i>1} \left( log\left(p_{ij}\right) - log\left(\hat{p}_{ij}\right) \right)^2, \tag{1}$$

where $N_{pairs} = (L-1) \times (L-2)/2$ is the number of residue pairs with sequence separation > 1 and $p_{ij}$ and $\hat{p}_{ij}$ are the contact frequencies for residues $i$ and $j$ in the reference and generated ensembles respectively. To avoid frequencies equal to zero, we use a pseudo-count value of 0.01. Contacts are defined with a Cα distance threshold of 8.0 Å.

In contrast to a similar score we used previously, this score only considers pairs of residues with a sequence separation of at least 2, since there is very little variability in the distance between two adjacent Cα atoms and predicting contact frequencies would be trivial. This modification is included in the remaining scores considering a pair of residues (MSE_d, aKLD_d and aJSD_d).

**MSE_d**

To evaluate average Cα-Cα interatomic distance values, we use the following MSE score:

$$\text{MSE\_d} = \frac{1}{N_{pairs}} \sum_{j-i>1} \left( m_{ij} - \widehat{m}_{ij} \right)^2, \tag{2}$$

where $m_{ij}$ and $\widehat{m}_{ij}$ are the average distances between the Cα atoms of residue $i$ and $j$ in the reference and generated ensembles.

**KLD approximations**

To compare mono-dimensional distributions of continuous features, we employ an approximation of the Kullback-Leibler divergence (KLD) by binning the values of the features as we did similarly to previous work[3]. We first take the minimum and maximum value of a feature over the reference and generated ensembles and split the range in $N_{bins} = 50$ equally-spaced bins. We then approximate KLD as:

$$\text{KLD}(P \parallel Q) = \sum_{k}^{N_{bins}} P_k \frac{log\left(P_k\right)}{log\left(Q_k\right)}, \tag{3}$$

where $k$ is the index of a bin and $P_k$ and $Q_k$ are the frequencies calculated in bin $k$ (using a pseudo-count value of 0.001) for the generated and reference ensembles respectively.

**aKLD_d**

To compare pairwise Cα-Cα interatomic distance distributions, we use the aKLD_d score, which is computed as:

$$\text{aKLD\_d} = \frac{1}{N_{pairs}} \sum_{j-i>1} KLD\left(\widehat{M}_{ij} \parallel M_{ij}\right), \tag{4}$$

where $M_{ij}$ and $\widehat{M}_{ij}$ are the distributions of Cα-Cα distances between residue $i$ and $j$ in the reference and generated ensembles.

**aKLD_t**

To compare distribution of α torsion angles, we use the aKLD_t score, which is computed as:

$$\text{aKLD\_t} = \frac{1}{N_{torsion}} \sum_i \text{KLD}(\hat{A}_{i,i+1,i+2,i+3} \parallel A_{i,i+1,i+2,i+3}), \quad (5)$$

where $N_{torsion} = L - 3$ is the number α angles in a peptide and where $A_{i,i+1,i+2,i+3}$ and $\hat{A}_{i,i+1,i+2,i+3}$ are the distributions of α angles among residue $i$ and its next 3 residues in the reference and generated ensembles.

**KLD_r**

To compare Cα radius-of-gyration distributions, we use the the KLD_r score:

$$\text{KLD\_r} = \text{KLD}(\hat{R} \parallel R), \quad (6)$$

where $R$ and $\hat{R}$ are the radius-of-gyration distributions for the reference and generated ensembles.

**aJSD_d and aJSD_t**

The KLD values that we use in the scores above, are asymmetric, since swapping the reference and proposed distributions would result in a different KLD value. Therefore, KLD-based scores assume that there is a reference distribution. To compare pairs of distributions $P$ and $Q$ in which we do not assume any of them to be a reference, we use the symmetric Jensen-Shannon (JS) divergence which we compute via the KLD approximation above:

$$\text{JSD} = \frac{1}{2}\text{KLD}(P \parallel M) + \frac{1}{2}\text{KLD}(Q \parallel M), \quad (7)$$

where $M$ is a mixture distribution for which we compute frequencies as:

$$M_k = \frac{1}{2}P_k + \frac{1}{2}Q_k. \quad (8)$$

The aJSD_d and aJSD_t scores that we use to compare some ensembles in this work are calculated similarly to the aKLD_d and aKLD_t scores defined above, but instead use this JSD approximation.

#### Supplementary Figures

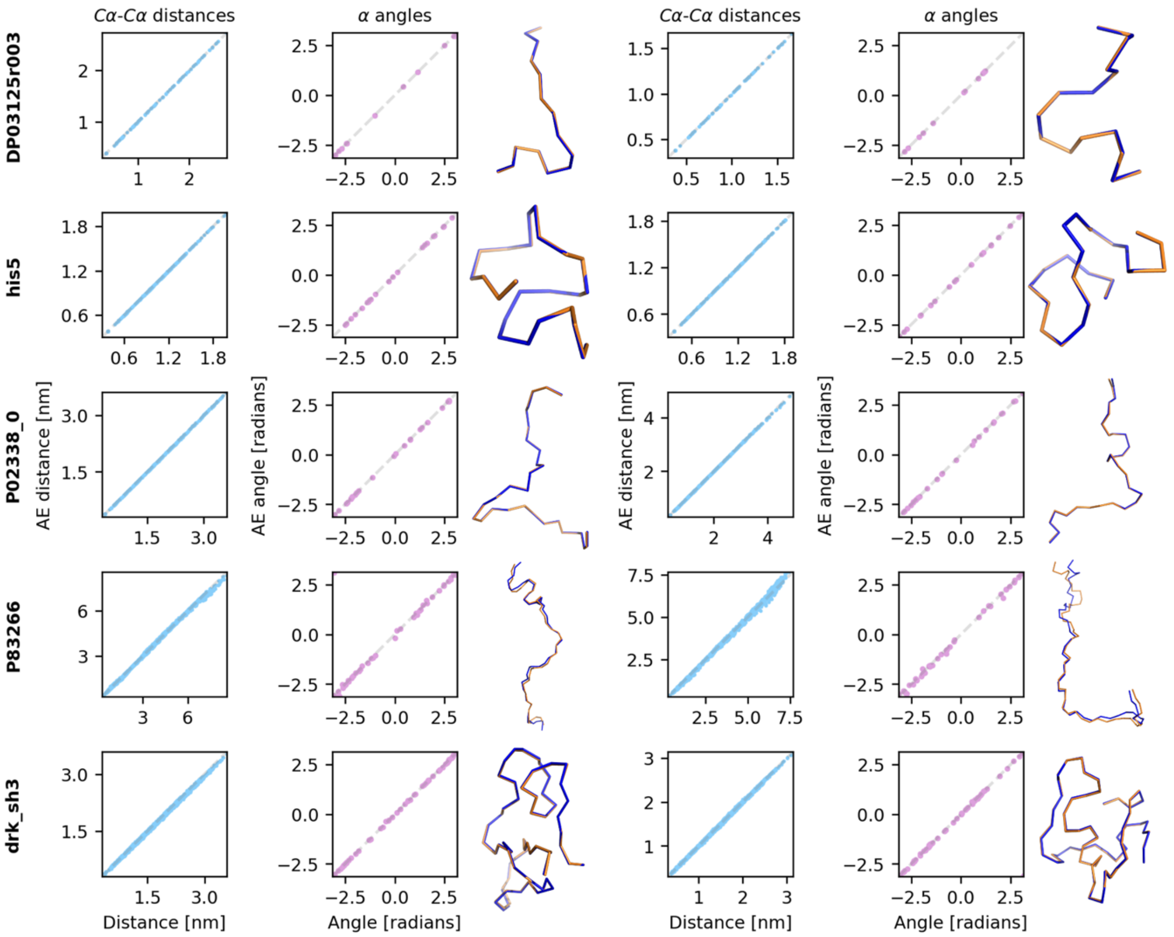


**S1 Fig. Cα conformations reconstructed by the AE.** Each row shows two MCMC conformations randomly extracted from simulation data of a test peptide (only five peptides are shown). Conformations were encoded and decoded back by the AE. For each conformation, we show: (i) on the left, a scatter plot with the original Cα-Cα distances against the corresponding values in the AE reconstruction; (ii) on the middle, a scatterplot with the original α angles against the reconstructed values; (iii) on the right, a superposition of the original (blue) and reconstructed (orange) Cα structures. In the scatterplots, original values are on horizontal axes, reconstructed values on vertical axes. Examples were randomly selected.

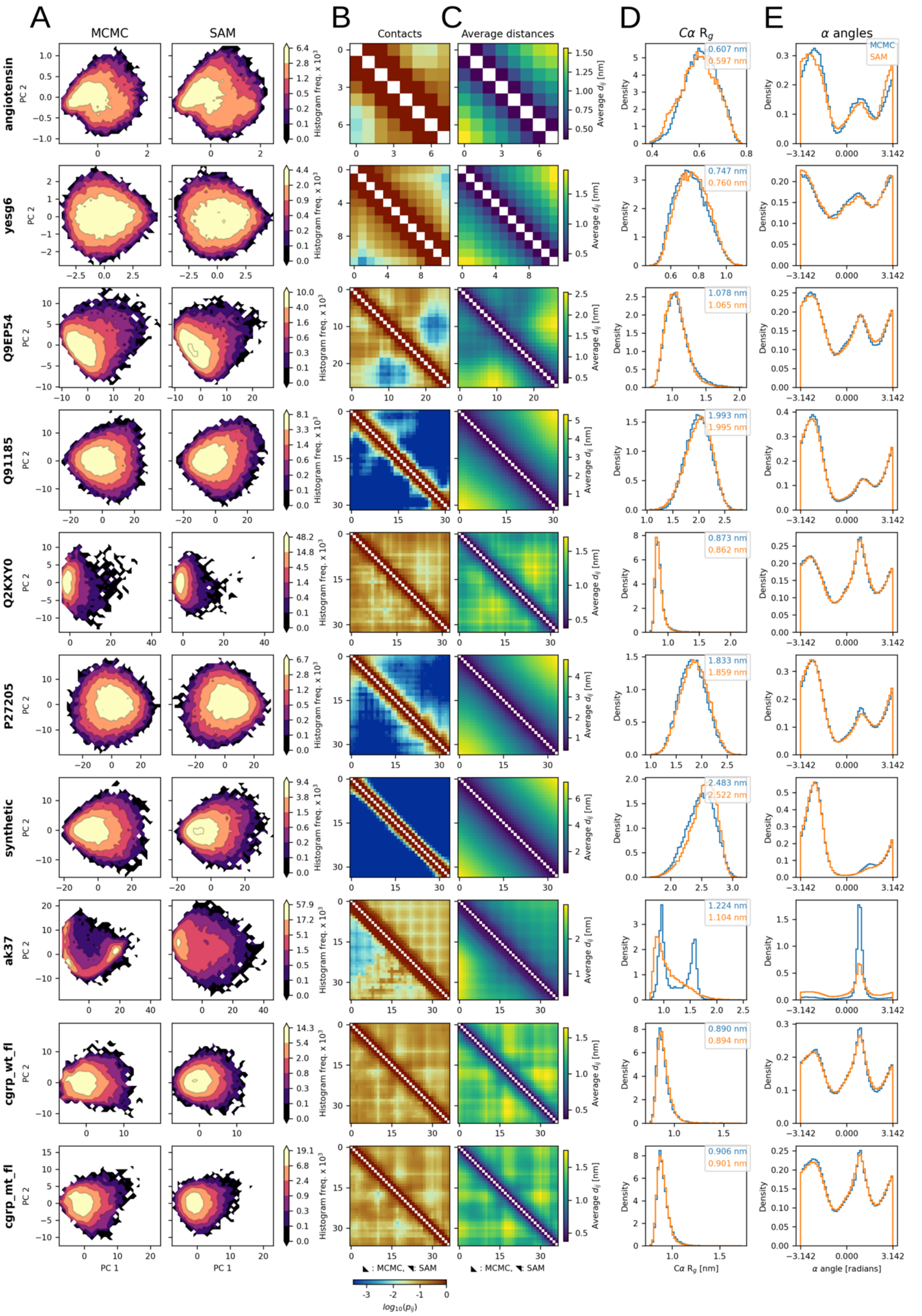


**S2 Fig. Additional structural ensembles from idpSAM.** Each row shows the ensembles of a peptide from the test set: angiotensin (*L* = 8), yesg6 (*L* = 12), Q9EP54 (*L* = 27), Q91185 (*L* = 32), Q2KXY0 (*L* = 33), P27205 (*L* = 34), synthetic (*L* = 34), ak37 (*L* = 37), cgrp_wt_fl (*L* = 37) and cgrp_mt_fl (*L* = 37). See **Fig 2** in the main text for more details.

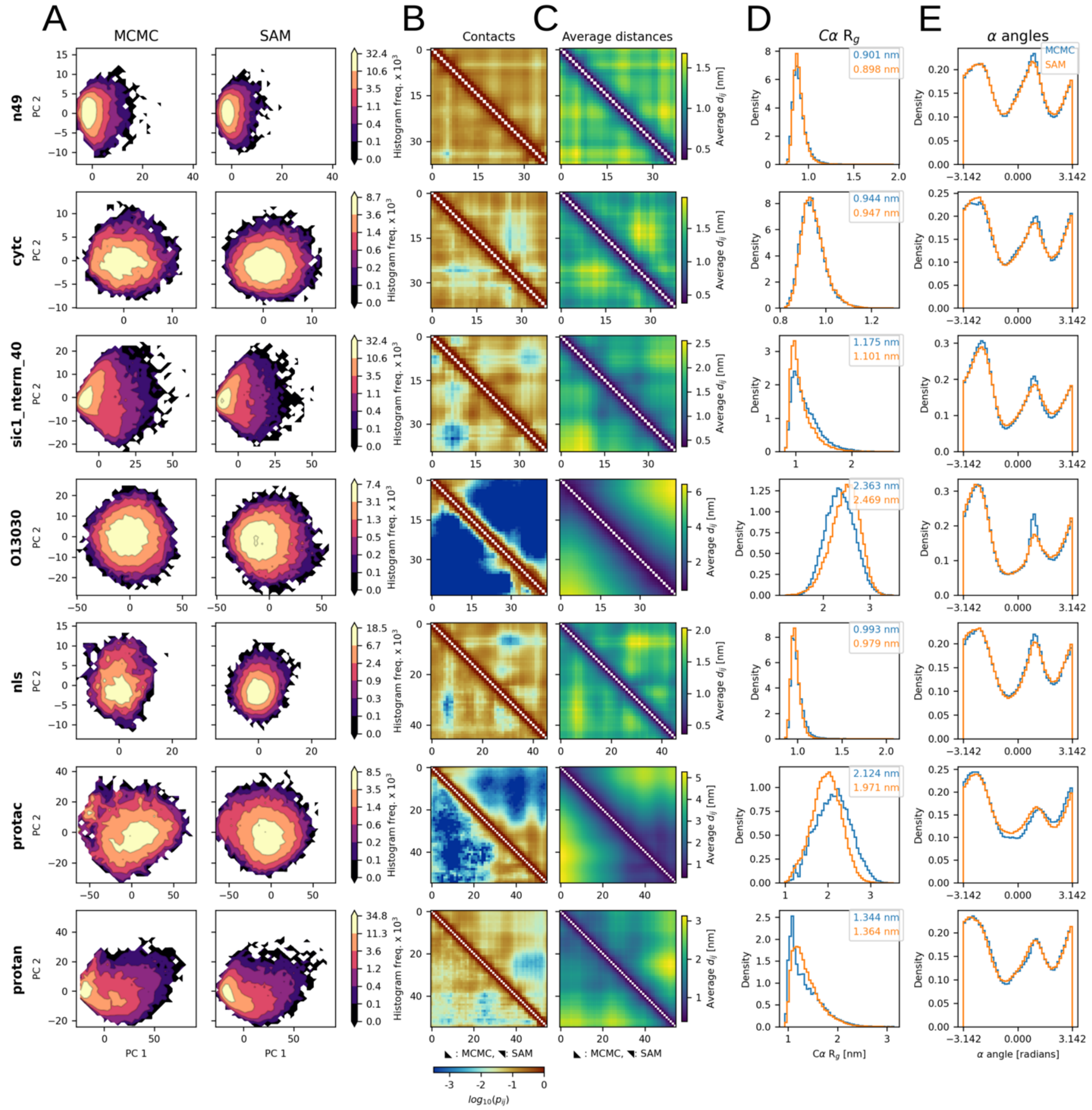

**S3 Fig. Additional structural ensembles from idpSAM.** Each row shows the ensembles of a peptide from the test set: n49 (*L* = 38), cytc (*L* = 39), sic1_nterm_40 (*L* = 40), O13030 (*L* = 44), nls (*L* = 46), protac (*L* = 55) and protan (*L* = 55). See **Fig 2** for more details.

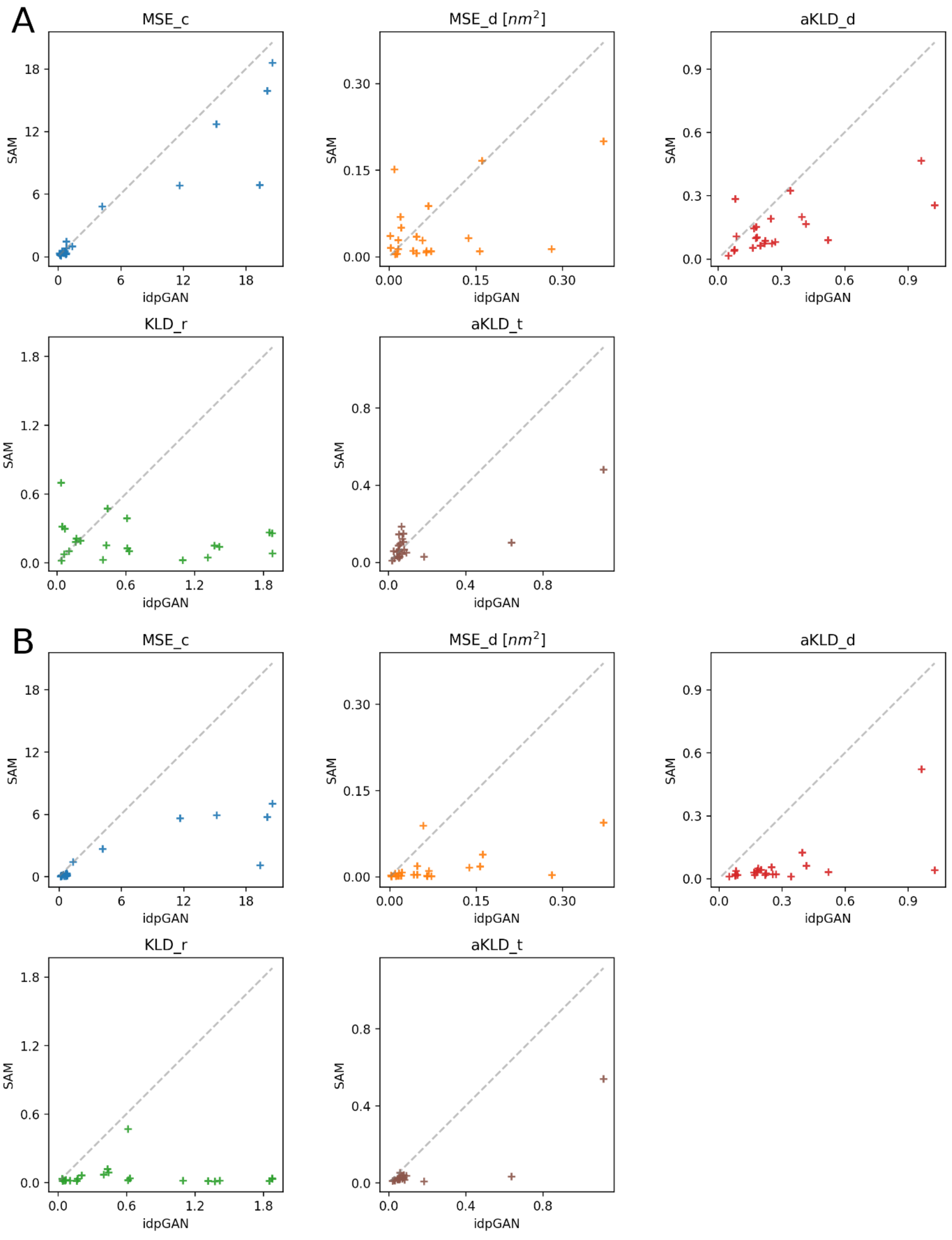


**S4 Fig. Evaluating SAM and idpGAN.** (A) Each subplot confronts the evaluation scores of the two models for the 22 test set peptides. The SAM version evaluated here was trained with the same dataset of idpGAN. (B) Similar confrontations, but the SAM version evaluated here was trained with the entire training set presented in this study.

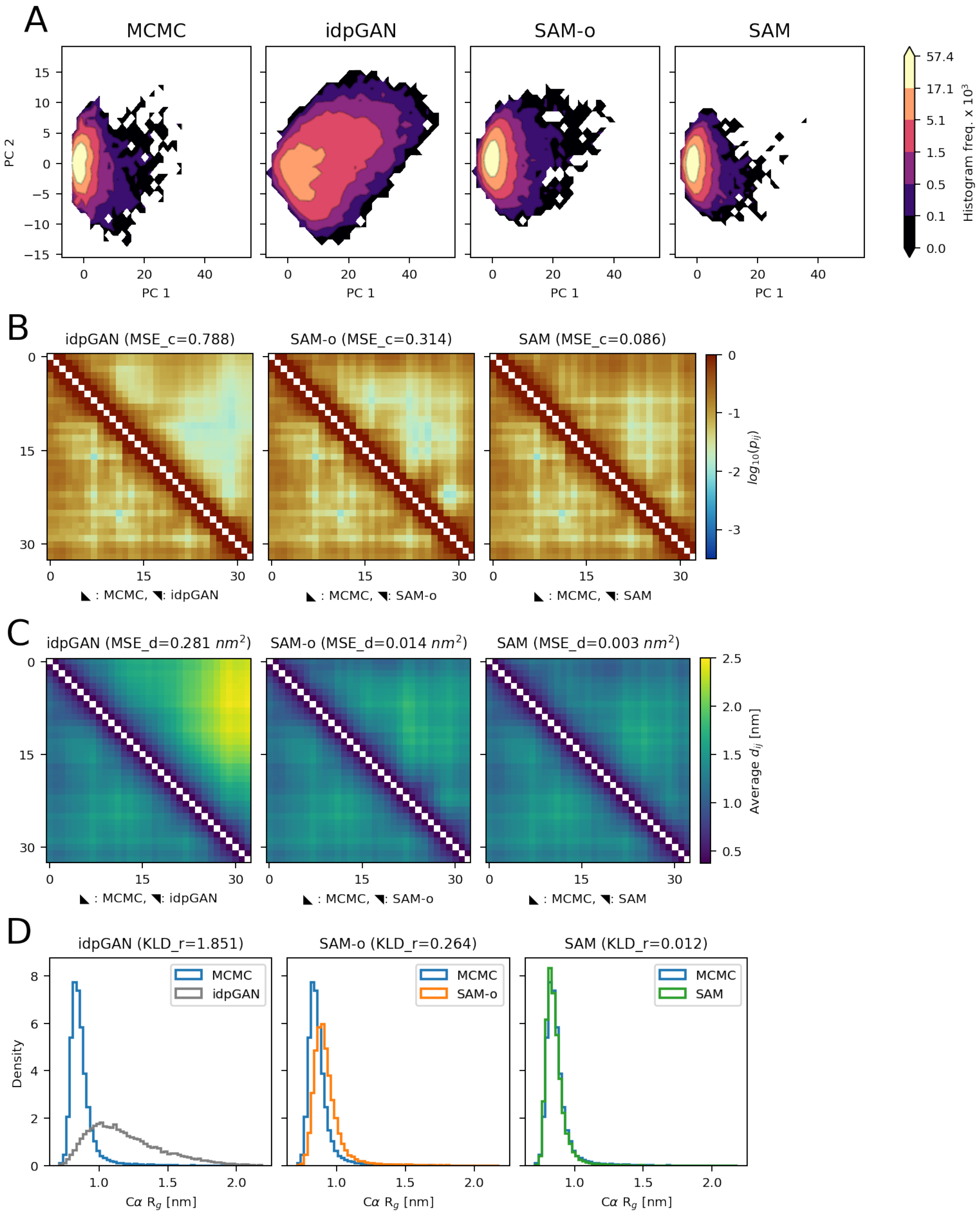


**S5 Fig. Modeling the ensemble of Q2KXY0 with SAM and idpGAN.** (A) PCA histograms for the structural ensembles of idpGAN, SAM-o and SAM-e. Here, SAM-o is a model trained with original training set of idpGAN (1089 IDRs), while SAM-e was trained with the expanded training set of this study (3,259 peptides) and is the model discussed in most of the main text. Frequency values in colorbars are multiplied by $1 \times 10^3$. (B) Cα-Cα contact maps of the three methods. The MSE_c scores of the ensembles are reported in brackets. (C) Average Cα-Cα distance maps of the three methods, with their MSE_d scores reported in brackets. (D) Cα $R_g$ histograms the three methods, with KLD_r values in brackets.

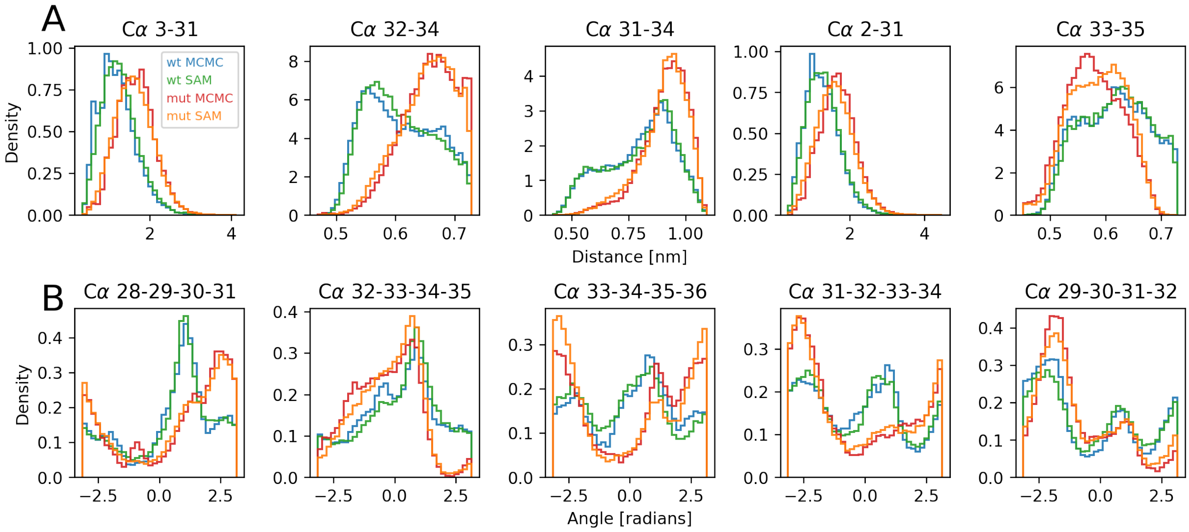


**S6 Fig. Cα-Cα distances and α torsion angle distributions in CGRP variants.** (A) Histograms of the five Cα-Cα distance distributions with the highest JS divergence between the wild-type and mutant ensembles from MCMC simulations. Residue indices of the Cα atoms are reported on top of the subplots (B). Histograms of the five α angle distributions with the highest JS divergence between the wild-type and mutant ensembles from MCMC. Residue indices of the Cα atoms defining the torsion angles are reported on top.

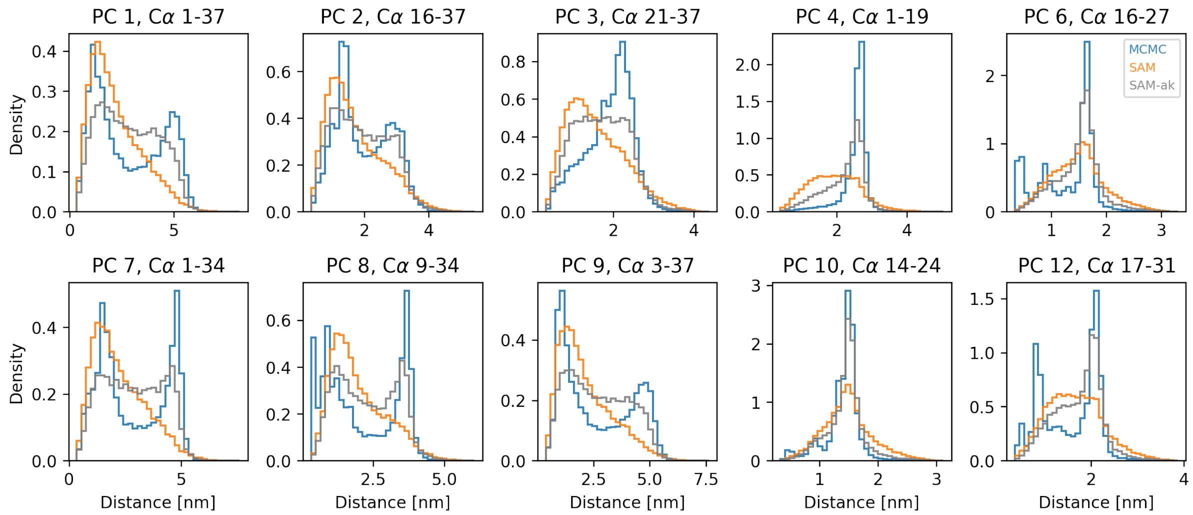


**S7 Fig. Cα-Cα distance distributions in ak37 ensembles.** The plots show data for those Cα-Cα distances that exhibit the highest absolute loading value for 10 of the top principal components (PCs) identified in a PCA on the MCMC ensemble (refer to the main text for PCA details). The distances associated with PC 5 and 11 are not shown since it is the same shown for PC 1. SAM-ak was trained on the full training set of SAM and additional simulation data from three peptides with sequences similar to ak37.

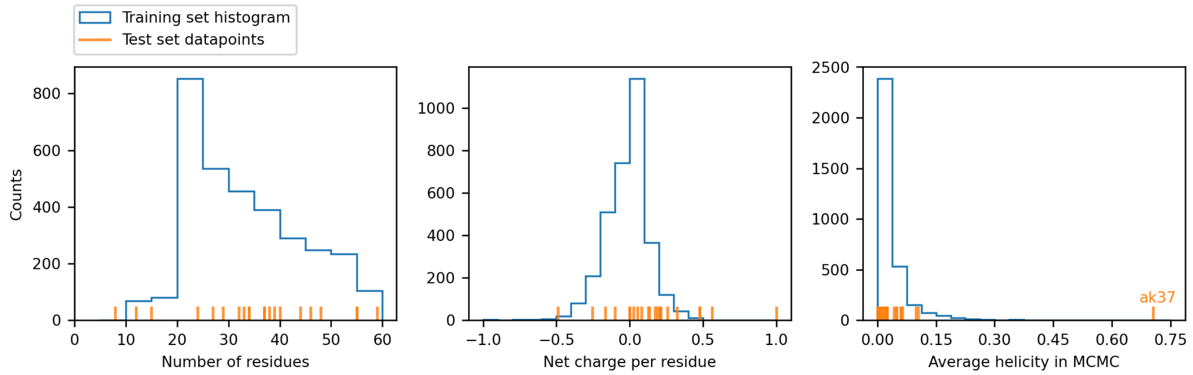


**S8 Fig. Properties of the 3,259 training and 22 test sequences.** Left panel: lengths of the peptides. Central panel: net charge per residue. Right panel: average helicity in an ensemble of 10,000 conformations from MCMC simulations. Helicity is defined as the fraction of residues in an all-atom peptide conformation found in a helical state according to the DSSP algorithm[18]. The average helicity value of the ak37 peptide is highlighted in the plot.

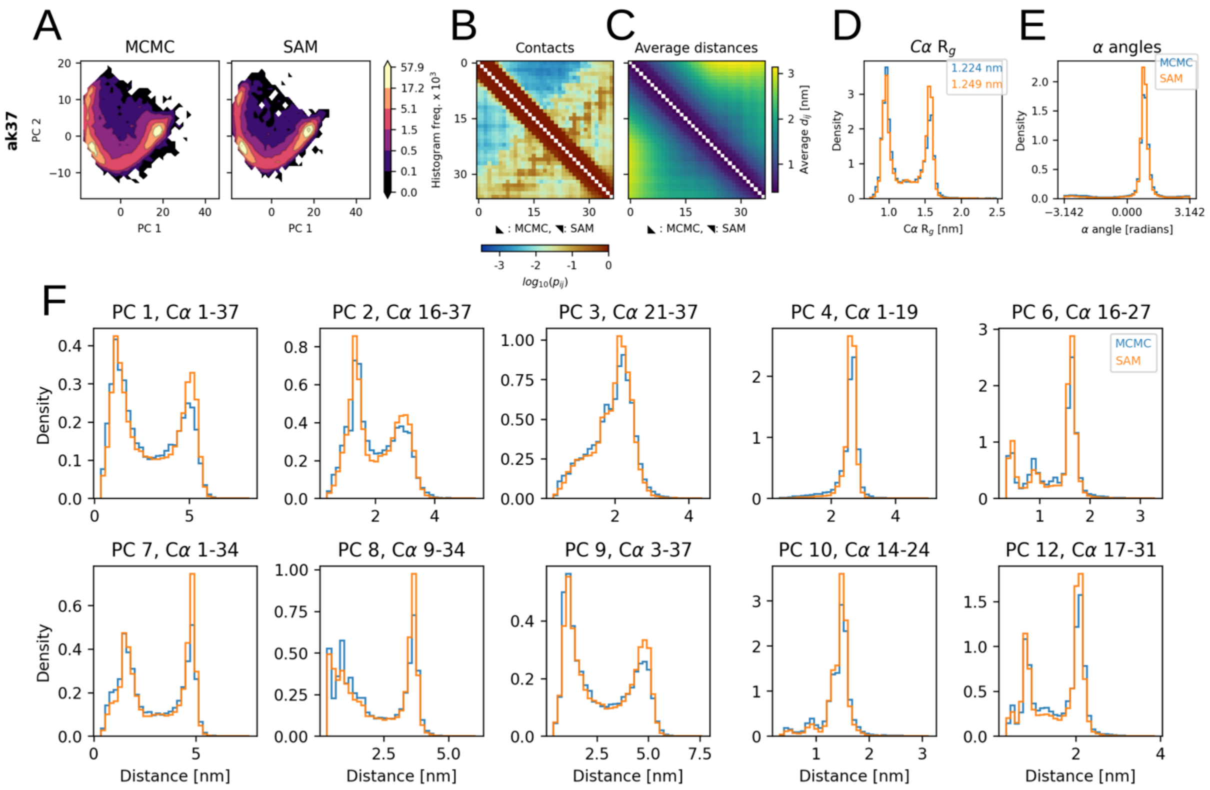

**S9 Fig. Overfitting on the structural ensemble of ak37.** (A) to (E) Ensemble of ak37 modeled by a SAM version trained only on ak37 data. See **Fig 2** in the main text for more details. (F) Cα-Cα distance distributions of the same ensemble. See **S7 Fig** for details.

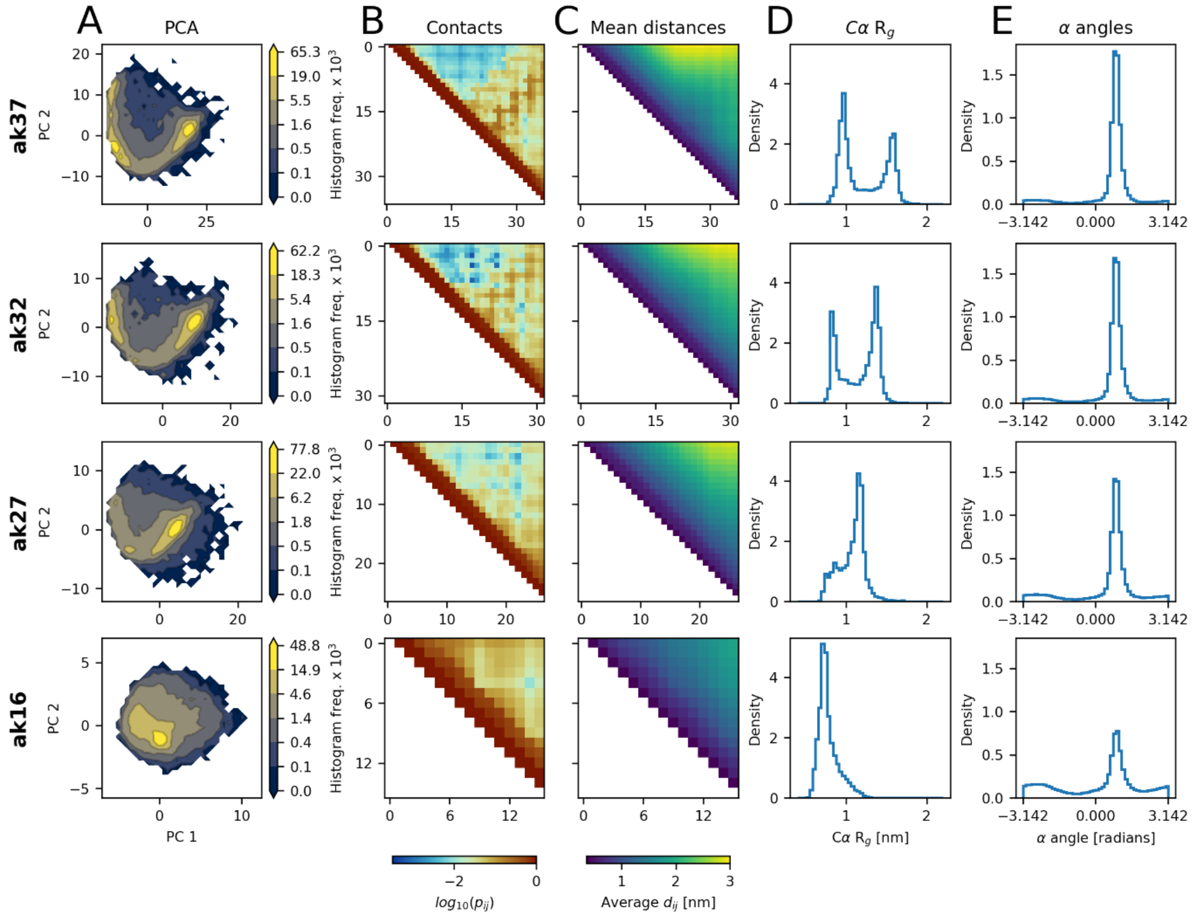


**S10 Fig. MCMC ensembles for the ak synthetic peptides.** The ensembles consist of conformations randomly extracted from MCMC simulations. (A) PCA histograms for the four peptides. PCA was performed on each peptide independently and each row uses its own principal axes. (B) Cα-Cα contact maps. (C) Average Cα-Cα distances. (D) Cα $R_g$ histograms. (E) α torsion angle histograms.

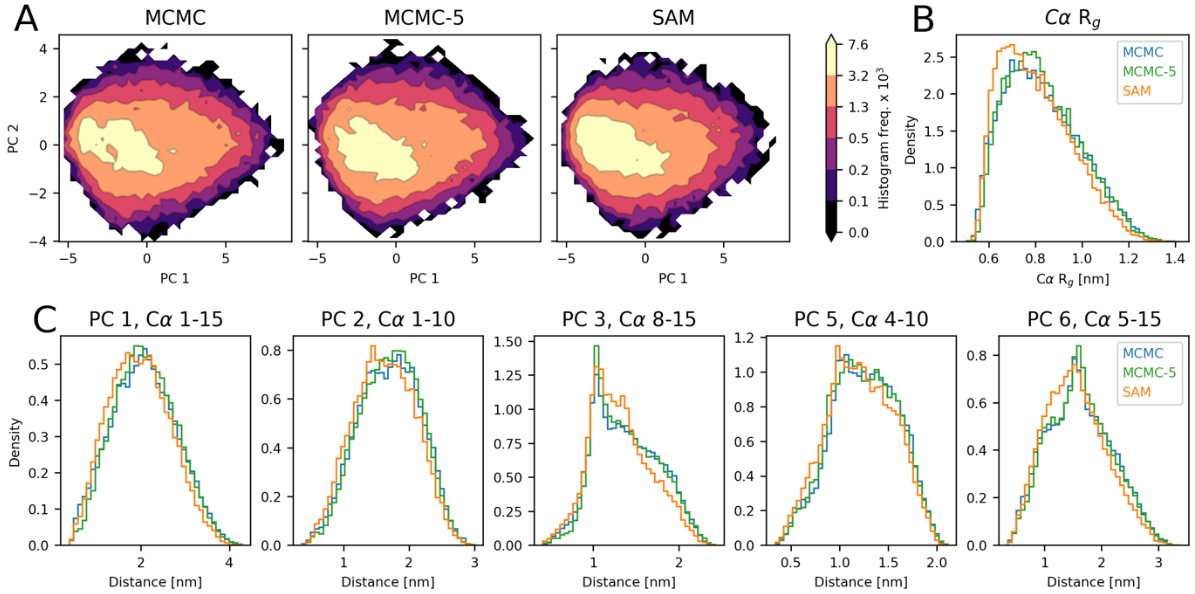


**S11 Fig. Modeling of the DP03125r003 peptide (*L* = 15) with SAM and different levels of MCMC sampling.** Refer to **Fig 5** in the main text for a description of the panels. The ensemble from only 5 MCMC runs closely approximates the one from extensive sampling consisting of 73 MCMC runs (**S2 Table**).

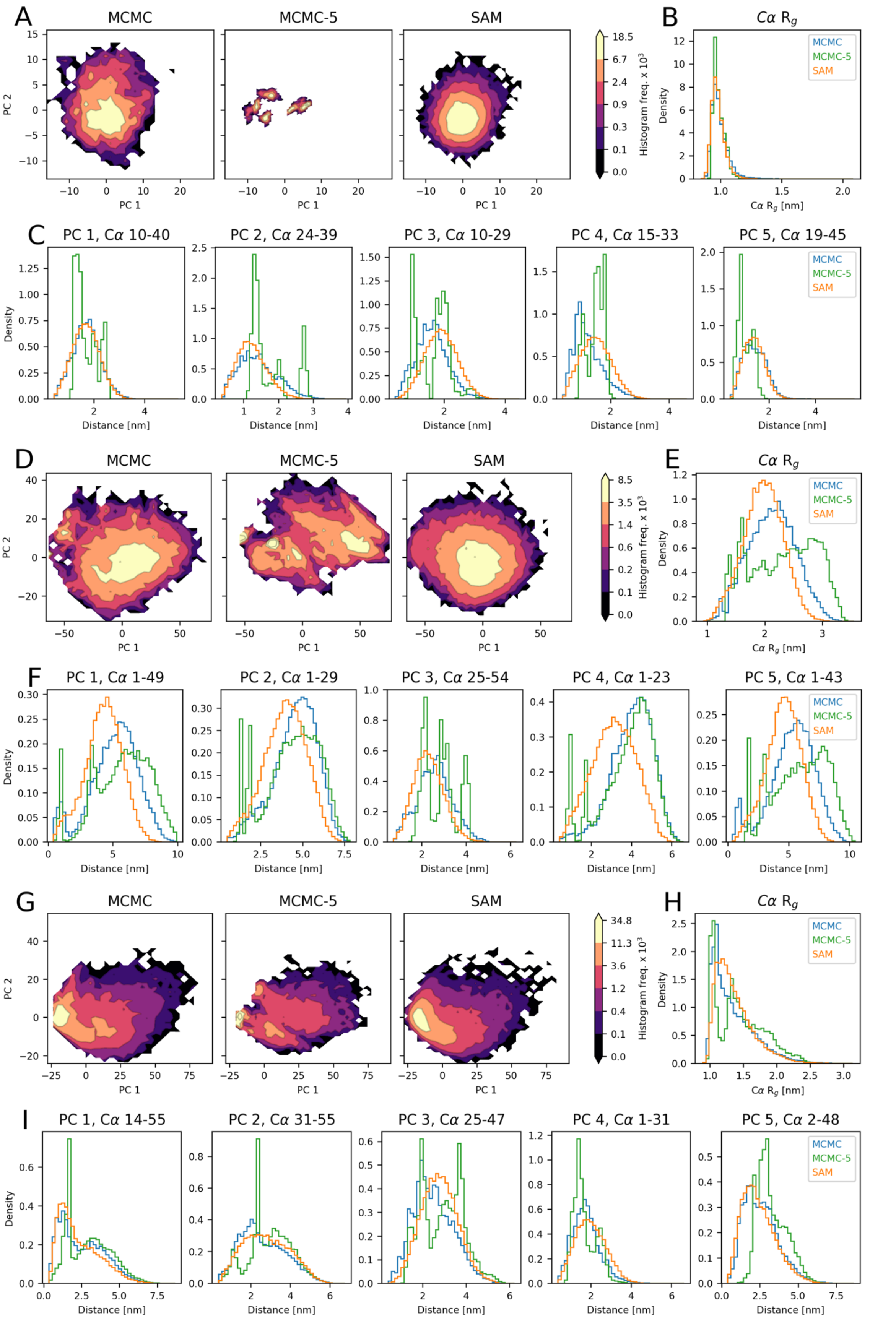


**S12 Fig. Modeling of the nls ($L$ = 46), protac ($L$ = 55) and protan ($L$ = 55) peptides with SAM and different levels of MCMC sampling.** Refer to **Fig 5** in the main text for a description of the panels. (A) to (C): data or nls. (D) to (F): data for protac. (G) to (I): data for protan.

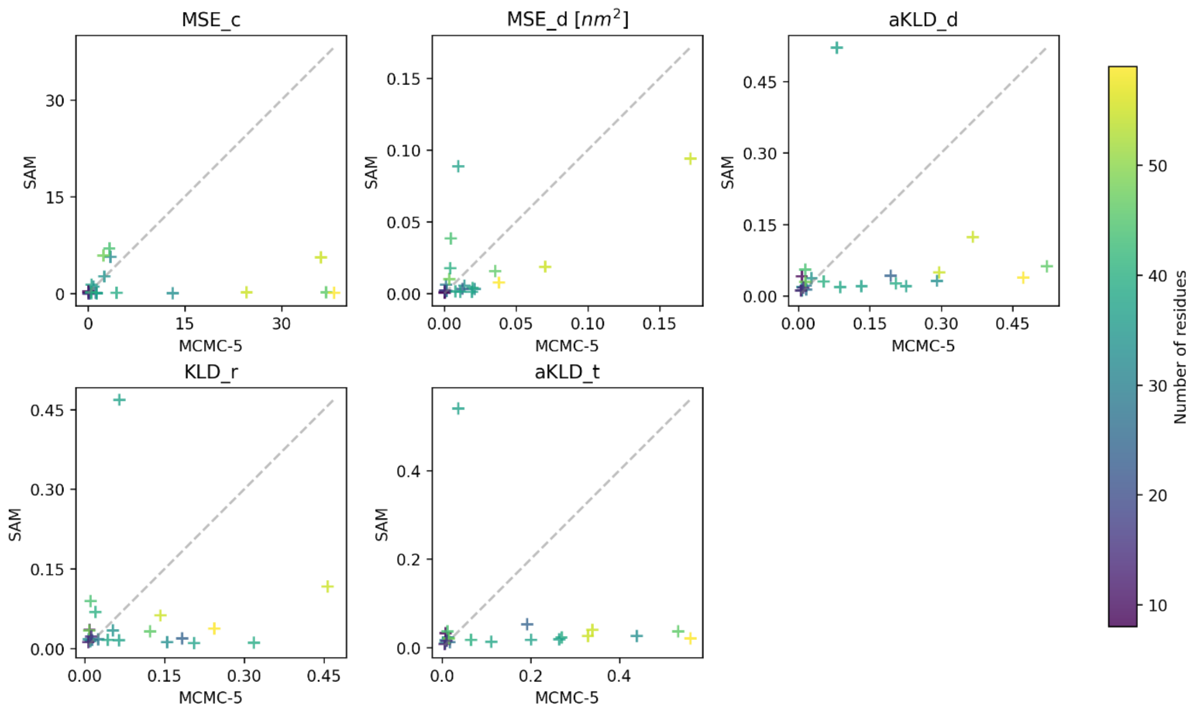


**S13 Fig. Evaluating SAM and the MCMC-5 sampling strategy.** Each subplot confronts the evaluation scores of the two strategies for the 22 test set peptides. Markers are colored according to the length of the corresponding peptide.

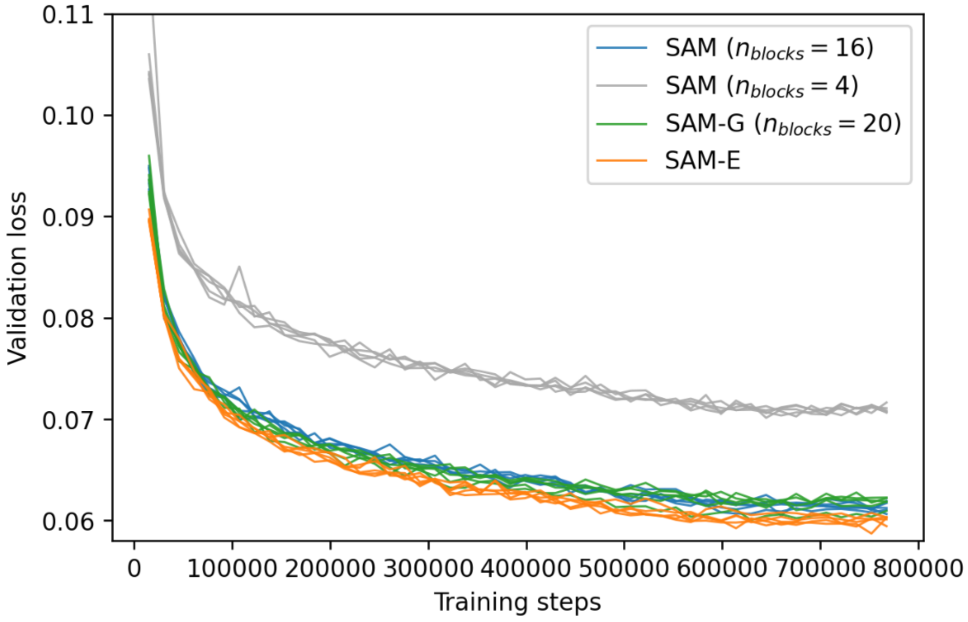


**S14 Fig. Validation losses observed during DDPM training for different idpSAM versions.** All the models with the "SAM" label have the same architecture for the noise prediction network, but have different number of transformer blocks in it. The SAM model with 20 blocks corresponds to the SAM-G model discussed in the main text. The SAM-E model has a different architecture, it has 16 blocks and incorporates 4 FrameDiff edge update operations. In case of the SAM models with 4 and 16 blocks, there is a large difference in validation loss at the end of training, which translates into a large difference in ensemble modeling performance (**S4 Table**). SAM-E has slightly better validation loss with respect to SAM versions with comparable numbers of layers, but it does not seem to significantly improve modeling performance (**Table 1**). For each model, we show validation loss curves from 5 different training runs.

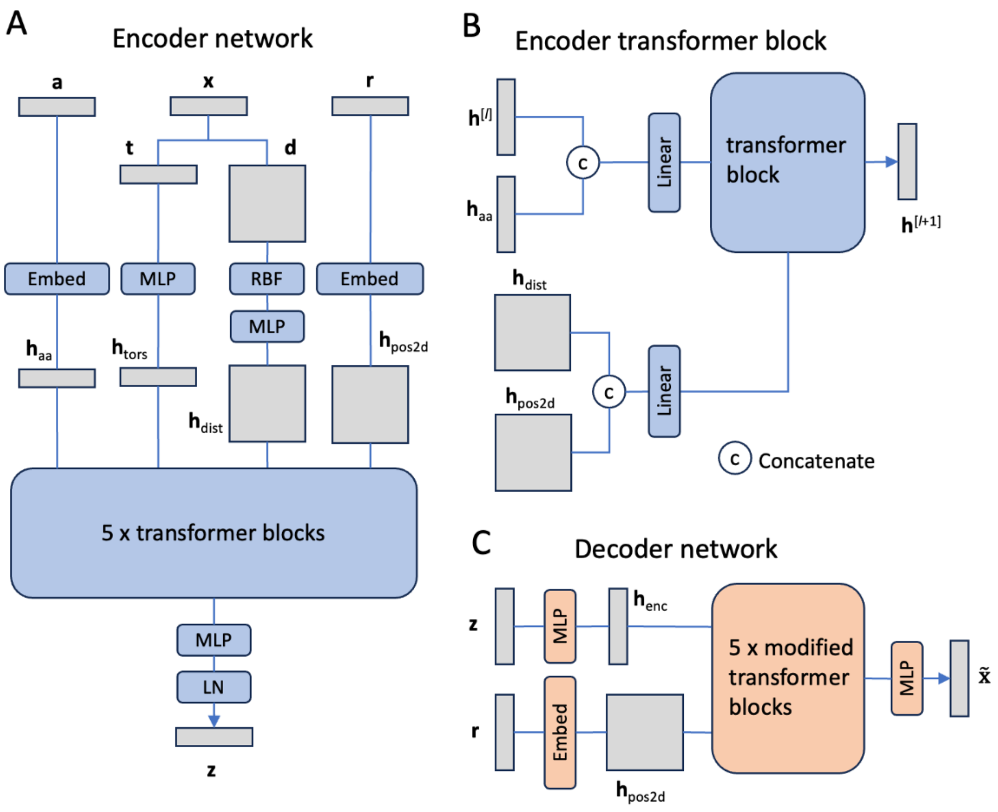


**S15 Fig. Encoder and decoder networks of SAM.** (A) Outline of the encoder network. The elements of the network are colored in blue, the tensors being processed by it are colored in gray. The tensor **r** represents the numerical indices of the input peptide, which are used to generate a 2d relative positional embedding $\mathbf{h}_{2dpos}$. Embed: embedding layer. MLP: multilayer perceptron. RBF: radial base function for embedding interatomic distances. LN: layer normalization. (B) Preparation of the input of a transformer block in the encoder. (C) Outline of the decoder network.

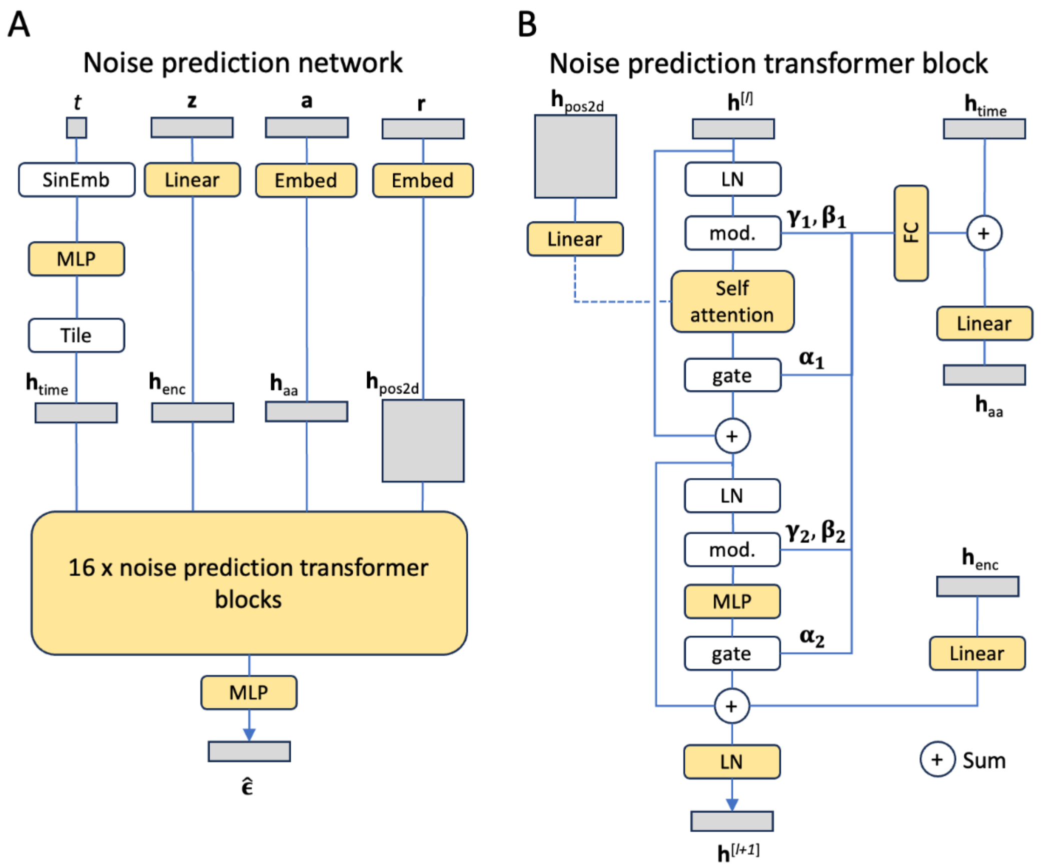


**S16 Fig. Noise prediction network of SAM.** (A) Outline of the entire noise prediction network. SinEmb: sinusoidal embedding. (B) Illustration of a transformer block of the network. FC: fully-connected module consisting of an activation and a linear layer. LN: layer normalization (colored in white if does not have learnable elementwise affine parameters). Mod.: modulate operation for adaLN-Zero. Gate: gate operation of adaLN-Zero.

### Supplementary Tables

**S1 Table. Reconstruction quality by the AE.**

| **Strategy** | $C_{dist}$ | $C_{tors}$ |
|---|---|---|
| AE[a] ($c = 16$) | (3.64 ± 1.33) × $10^{-3}$ | (1.96 ± 0.69) × $10^{-3}$ |
| AE ($c = 4$) | (4.13 ± 0.73) × $10^{-2}$ | (6.22 ± 1.06) × $10^{-3}$ |
| AE ($c = 8$) | (7.34 ± 2.63) × $10^{-3}$ | (3.10 ± 0.86) × $10^{-3}$ |
| AE ($c = 32$) | (2.49 ± 0.89) × $10^{-3}$ | (1.63 ± 0.59) × $10^{-3}$ |
| Perturbed[b] (0.010 Å) | (6.91 ± 1.03) × $10^{-4}$ | (5.90 ± 0.10) × $10^{-5}$ |
| Perturbed (0.025 Å) | (4.30 ± 0.64) × $10^{-3}$ | (3.70 ± 0.08) × $10^{-4}$ |
| Perturbed (0.05 Å) | (1.72 ± 0.26) × $10^{-2}$ | (1.48 ± 0.03) × $10^{-3}$ |
| Perturbed (0.1 Å) | (6.89 ± 1.02) × $10^{-2}$ | (5.95 ± 0.13) × $10^{-3}$ |

For each test set peptide, 10,000 snapshots were randomly extracted from MCMC data and average $C_{dist}$ and $C_{tors}$ values were computed between these snapshots and their autoencoded or perturbed versions. The table reports the mean (with standard errors) of these average values for the 22 test peptides.

[a]MCMC conformations were encoded and decoded by AE of idpSAM. The AE of the default idpSAM version has an encoding dimension $c = 16$.

[c]MCMC conformations in which Cα positions were perturbed by adding different levels of Gaussian random noise (standard deviations are reported in parentheses).

**S2 Table**. **Properties of the 22 test set peptides.**

| Name | Sequence[a] | L[b] | q/L[c] | UniProt[d] | 298 K runs[e] | RE runs[f] |
|---|---|---|---|---|---|---|
| angiotensin | DRVYIHPF | 8 | 0.00 | P01019 | 80 | - |
| yesg6 | YESGGGGGGATD | 12 | -0.17 | - | 90 | - |
| DP03125r003 | KPSNCQNKESASKQS | 15 | 0.13 | O14965 | 73 | - |
| his5 | DSHAKRHHGYKRKFHEKHHSHRGY | 24 | 0.21 | P15516 | 149 | - |
| Q9EP54 | MACYPVNIRARGLGKNMGMKSRGRGKG | 27 | 0.26 | **Q9EP54** | 150 | - |
| P02338_0 | MRSFDQGSTRAPARERCRRQRPEGRSAQR | 29 | 0.21 | **P02338** | 150 | - |
| Q91185 | MRRQASLPARRRRRVRRTRVVRRRRRVGRRRH | 32 | 0.56 | **Q91185** | 75 | - |
| Q2KXY0 | MFDNASTRNNKRERGKRQGKQTRTQRHADRSQT | 33 | 0.21 | **Q2KXY0** | 175 | - |
| P27205 | AGSKSRSRSRSRSRSKSPAKSASPKSAASPRASR | 34 | 0.32 | **P27205** | 150 | - |
| synthetic | RRRRRRRRRRRRRRRRRRRRRRRRRRRRRRRRRR | 34 | 1.00 | - | 60 | - |
| ak37 | AAKAAAAKAAAAKAAAAKAAAAKAAAAKAAAAKAAGY | 37 | 0.19 | - | 149 | - |
| cgrp_wt_fl | ACDTATCVTHRLAGLLSRSGGVVKNNFVPTNVGSKAF | 37 | 0.08 | **P06881**[*] | 175 | - |
| cgrp_mt_fl | ACDTATCVTHRLAGLLSRSGGVVKNNFVPTDVGPWSF | 37 | 0.03 | **P06881**[*] | 169 | - |
| n49 | GCQTSRGLFGNNNTNNINNSSSGMNNASAGLFGSKPFA | 38 | 0.05 | Q02199 | 194 | - |
| cytc_nter | MIFFMVMPIMIGGFGNWLVPLMIGAPDMAFPRMNNSFWL | 39 | 0.00 | P00395 | 184 | - |
| sic1_nterm_40 | MTPSTPPRSRGTRYLAQPSGNTSSSALMQGQKTPQKPSQN | 40 | 0.12 | P38634 | 158 | - |
| O13030 | MAYGRARSRGRSVRRRRRGRSPGRRRRGRRSDNDAPRRRRRRRQ | 44 | 0.48 | **O13030** | 75 | - |
| nls | ACETNKRKREQISTDNEAKMQIQEEKSPKKKRKKRSSKANKPPEFA | 46 | 0.17 | Q03281 | 184 | 69 |
| P83266 | ARRRHSMKKKRKSVRRRKTRKNQRKRKNSLGRSFKQHGFLKQPPRFRP | 48 | 0.48 | **P83266** | 65 | - |
| protac | CEEGGEEEEEEEEGDGEEEDGDEDEEAESATGKRAAEDDEDDDVDTKK QKTDEDC | 55 | -0.49 | P06454-2 | 137 | 56 |
| protan | CDAAVDTSSEITTKDLKEKKEVVEEAENGRDAPANGNANEENGEQEAD NEVDEEC | 55 | -0.25 | P06454-2 | 137 | 59 |
| drk_sh3 | MEAIAKHDFSATADDELSFRKTQILKILNMEDDSNWYRAELDGKEGLI PSNYIEMKNHD | 59 | -0.10 | Q08012 | 166 | 69 |

[a]Positively charged residues are in blue, negatively charged in red.

[b]Number of residues in a peptide.

[c]Net charge per residue of a peptide.

[d]UniProt accession number of the sequence. Synthetic peptides have an empty value. Bold font corresponds to sequences covering their entire UniProt entry.

[e]Number of MCMC runs at 298 K.

[f]Number of replica exchange (RE) MCMC runs.

[*]Full sequence of the biologically-active peptide.

**S3 Table. Properties of the ak synthetic peptides.**

| Name | Sequence[a] | L[b] | q/L[c] | 298 K runs[d] |
|---|---|---|---|---|
| ak16 | YGCAKAAAAKACAAKA | 16 | 0.19 | 5 |
| ak27 | AAKAAAAKAAAAKAAAAKAAAAKAAGY | 27 | 0.19 | 5 |
| ak32 | AAKAAAAKAAAAKAAAAKAAAAKAAAAKAAGY | 32 | 0.19 | 5 |
| ak37 | AAKAAAAKAAAAKAAAAKAAAAKAAAAKAAAAKAAGY | 37 | 0.19 | 149 |

[a]Positively charged residues are in blue, negatively charged in red.
[b]Number of residues in a peptide.
[c]Net charge per residue of a peptide.
[d]Number of MCMC runs at 298 K.

**S4 Table. Evaluation scores of ensembles from modified versions of SAM.**

| Strategy | MSE_c | MSE_d [$nm^2$] | aKLD_d | KLD_r | aKLD_t |
|---|---|---|---|---|---|
| SAM[a] | 1.421 ± 0.499 | 0.015 ± 0.006 | 0.057 ± 0.023 | 0.053 ± 0.021 | 0.047 ± 0.024 |
| SAM-ak[b] | 1.470 ± 0.479 | 0.013 ± 0.005 | 0.044 ± 0.008 | 0.051 ± 0.013 | 0.029 ± 0.005 |
| $n_{blocks} = 4$[c] | 2.649 ± 1.061* | 0.024 ± 0.008* | 0.092 ± 0.030* | 0.094 ± 0.025* | 0.075 ± 0.037* |
| $n_{blocks} = 8$ | 1.786 ± 0.641 | 0.013 ± 0.003 | 0.067 ± 0.027* | 0.060 ± 0.018 | 0.060 ± 0.032* |
| no-lr-sched[d] | 2.326 ± 1.061* | 0.021 ± 0.008* | 0.065 ± 0.021* | 0.092 ± 0.037* | 0.049 ± 0.015* |
| no-adalnzero[e] | 3.926 ± 1.656 | 0.038 ± 0.015 | 0.093 ± 0.030 | 0.167 ± 0.064* | 0.056 ± 0.026 |
| no-input-inject[f] | 1.373 ± 0.449 | 0.018 ± 0.010 | 0.063 ± 0.028 | 0.077 ± 0.037 | 0.057 ± 0.032* |
| $c = 4$[g] | 11.582 ± 6.616* | 0.423 ± 0.384* | 0.740 ± 0.56* | 1.062 ± 0.621* | 0.131 ± 0.037* |
| $c = 8$ | 1.546 ± 0.464 | 0.015 ± 0.004 | 0.096 ± 0.035* | 0.138 ± 0.083 | 0.065 ± 0.016* |
| $c = 32$ | 1.779 ± 0.603 | 0.018 ± 0.007 | 0.074 ± 0.031 | 0.068 ± 0.028 | 0.067 ± 0.036* |

Average scores are reported along with standard errors for the 22 test set peptides. Unless specified, all models were trained with the full training set of the default SAM version. Unless specified, all models use the same AE with only the DDPM being re-trained.

[a]Default idpSAM version.

[b]IdpSAM version with a DDPM trained with the full training set and three ak synthetic peptides.

[c]IdpSAM version with 4 transformer blocks in the noise prediction network of the DDPM.

[d]A "linear with warm up" learning rate schedule is not used to train the DDPM.

[e]The adaLN-zero mechanism is not used in the transformer blocks of the noise prediction network. Instead, time step and amino acid embeddings are projected to the same dimension of the node embeddings and added to them before the first layer normalization operation of the block.

[f]The initial node embedding is not inject at every block of the noise prediction network (**S16 Fig**).

[g]IdpSAM version with encoding dimension $c = 4$. Both the AE and DDPM were re-trained.

[*]Asterisks denote a statistically significant difference (using a Wilcoxon signed-rank test with a significance level of 0.05) between the scores of a strategy and of the default idpSAM version.

**S5 Table. Properties of the 25 validation set peptides.**

| Name[a] | Sequence[b] | L[c] | q/L[d] |
|---|---|---|---|
| A0A0G2JEB6 (2015-2035) | MPDHMRSSDYAADQSSSHAE | 20 | -0.15 |
| A0A452G813 (22-44) | IGDQPNDSYCYNSAKNSTVLQG | 22 | -0.05 |
| Q54WT6 (92-120) | RLIISAFKEIKFENQEKTKLTKFEKYQK | 28 | 0.14 |
| C6JUP1 (37-66) | KDKCEKYAVPVMRGKFYFSYQCTSKCHEG | 29 | 0.10 |
| P45198 (5-34) | QCNMKWGAEEEKKIELQAQGLQPDTLFSD | 29 | -0.10 |
| Q7NPQ7 (3-34) | QPSVKKICRNCKIIRRNRVVRVICTDPRHKQ | 31 | 0.29 |
| Q924W7 (711-744) | LPEVSYQFPKLDRPTKQMREAEERLKAIPQFCF | 33 | 0.03 |
| P54944 (19-54) | HYEYNPYPQQDVYDPYQMDRQPALERRIAALERQN | 35 | -0.06 |
| P72021 (34-72) | VIYHSQGLNALQMQNQDKNLEIRNDEYLELAKEFFDKA | 38 | -0.16 |
| P49525 (107-145) | ELVRAADMDAIEDDKVTIVGPDLKDMEEGKTYPWAMIF | 38 | -0.08 |
| A0FKN6 (23-62) | SNQDPIVDGMRLVEGDMLFDDGPLFTERNAVKYDQQLWP | 39 | -0.13 |
| B2GB76 (161-200) | NAVNLTDGLDGLVTGLATISFAAYLVLALVQGQTEVALF | 39 | -0.08 |
| A6W5T3 (77-117) | RRVGGSTYQVPIEVRPTRSTTLALRWLVGYARQRREKTMT | 40 | 0.18 |
| B8IMM7 (3-43) | TYQPSKLVRKRRHGFRARMATVGGRRVIAARRARGRKRLS | 40 | 0.38 |
| P30071 (285-326) | QKLSKSSRAWMREHLDDPFVKKAQKEGYRARAAYKLLEIQE | 41 | 0.02 |
| B9L6M9 (2-43) | RSLKKGPFVDDHLMKKVLKAKEEKNPKPIKTWSRRSTITPD | 41 | 0.17 |
| A3M851 (7-48) | ETPFISNNTIKKVNILVPIILSISDKEKFFNVLGHPTSNLF | 41 | 0.10 |
| A1S9H3 (305-347) | WEGDVYTMSAATREGTKELAEKLFDFIKSLPDEAAAADPDKE | 42 | -0.14 |
| Q9BKZ9 (61-103) | FKEAYEEEESREYVIENGVKRLVNQHYDSRGYSSAGRGQKGR | 42 | 0.00 |
| P70441 (91-134) | DPETDERLKKLGVSIREELLRPQEKSEQAEPPAAADTHEAGDQ | 43 | -0.14 |
| Q6IEU7 (245-289) | TIVTLFYGTLFCMYVRPPSEKSVEESKIIAVFYTFLSPMLNPLI | 44 | 0.00 |
| Q99PJ7 (274-318) | LLAIHQRTHTGEKPYTCLECSRRFRQRTALVIHQRIHTGEKPYP | 44 | 0.11 |
| B7IJW2 (1-48) | ELVTIEKGNIFFKSLEYKDLEEELLKKVSLYPNLEEYSKEEILTIET | 47 | -0.09 |
| A6UQD9 (114-161) | SFNLRGAVLANVSGNSQDQLQETIVDAIQSGEEKMLPGLGVLFEVIW | 47 | -0.13 |
| Q53RK8 (70-120) | SSDRDRDRDRDRDRDRDRDRERRRDDDSRRDRDRDRDRGDSSRRDR | 50 | 0.04 |

For all sequences, 3 MCMC runs were performed at 298K.

[a]UniProt accession number of the sequence with the starting and ending residue positions in the full sequence shown in parentheses.

[b]Positively charged residues are in blue, negatively charged residue in red.

[c]Number of residues.

[d]Net charge per residue.

**S6 Table. Details of SAM neural network training.**

| **Training process feature** | **Autoencoder** | **Diffusion model** |
|---|---|---|
| $n_{systems}$[a] | 1,239 ("part 1" and "part 2" of the full training set, see **S1 Text**) | 3,259 (full training set) |
| $n_{frames}$[b] | 1,580 | 300 |
| Number of training epochs | 60 | 50 |
| Batch size | 64 | 64 |
| Number of training steps | ~1,838,000 | ~765,000 |
| Optimizer[c] | Adam | Adam |
| Learning rate schedule | Milestone rate. Starts with lr=0.0005, multiplies the lr by 0.5 at epoch 10, 20, 30, 40 and 50. | Linear with warm up. Starts at lr=0.0, linearly warms up to lr=0.0005 in the first 10,000 steps, then linearly drops reaching lr=$5.0\times10^{-6}$ after another 615,000 steps. Stays at the same value until the rest of training. |
| Time required for training[d] | 1-2 days | 2-3 days |

[a]Number of different peptides used in the training set.
[b]Number of MCMC snapshots extracted from the simulation data of a single peptide during a training epoch.
[c]With the exception of the initial learning rate, we kept all other Adam parameters at their default PyTorch value.
[d]On a single NVIDIA RTX2080Ti GPU, using PyTorch 1.11.0 with automatic mixed precision training.
